# A Machine Learning Classifier to Identify and Prioritise Genes Associated with Cardiac Development

**DOI:** 10.1101/2024.11.08.622603

**Authors:** Mitra Kabir, Verity Hartill, Gist H. Farr, Wasay Mohiuddin Shaikh Qureshi, Stephanie L. Baross, Andrew J. Doig, David Talavera, Bernard D. Keavney, Lisa Maves, Colin A. Johnson, Kathryn E. Hentges

## Abstract

Congenital heart disease (CHD) is a major cause of infant mortality and presents life-long challenges to individuals living with these conditions. Genetic causes are known for only a minority of types of CHD. Discovering further genetic causes is limited by challenges in prioritising candidate CHD genes. We examined a wide range of features of mouse genes, including sequence characteristics, protein localisation and interaction data, developmental expression data and gene ontology annotations. Many features differ between cardiac development and non-cardiac genes, suggesting that these two gene types can be distinguished by their attributes. Therefore, we developed a supervised machine learning (ML) method to identify *Mus musculus* genes with a high probability of being involved in cardiac development. These genes, when mutated, are candidates for causing human CHD. Our classifier showed a cross-validation accuracy of 81% in detecting cardiac and non-cardiac genes. From our classifier we generated predictions of the cardiac development association status for all protein-coding genes in the mouse genome. We also cross-referenced our predictions with datasets of known human CHD genes, determining which are orthologues of predicted mouse cardiac genes. Our predicted cardiac genes have a high overlap with human CHD genes. Thus, our predictions could inform the prioritisation of genes when evaluating CHD patient sequence data for genetic diagnosis. Knowledge of cardiac developmental genes may speed up reaching a genetic diagnosis for patients born with CHD.

**Author Summary:** Congenital heart disease arises during pregnancy when the heart has formed incorrectly. These malformations affect ∼1% of newborns. Yet, despite their frequency, the underlying causes are still not known in many cases. It is clear that genetic factors contribute to these defects, and increasingly DNA sequencing is used to attempt to determine if an individual has a genetic change causative of their condition. However, analysis of patient sequence data often reveals changes that are difficult to interpret due to a lack of knowledge of the function of the gene harbouring a sequence change. We aimed to facilitate the process of sequence evaluation by predicting which genes of unknown function are likely involved in heart formation. Our predictions agree with novel experimental evidence about genes needed for heart development. We found that when mutated, a high proportion of the predicted cardiac genes do indeed cause heart defects. This result suggests that our predictions may be informative for expanding our understanding of the genetic basis of congenital heart disease.

## Introduction

Congenital heart disease (CHD), resulting from errors in the formation of the heart during gestation, is the most common structural malformation present at birth, affecting approximately 1.35 million infants annually worldwide [1], and is the primary contributor to infant mortality arising from birth defects [2]. CHD includes a wide variety of cardiac and great vessel malformations with varying severity in phenotype [3]. Advances in detection and surgical interventions have led to increased survival rates among children with CHD; however, patients seem to develop a wide range of co-morbidities. For example, psychological illness persists as a significant co-morbidity in adolescents and adults affected by the condition [4]. Children with CHD have a higher rate of neurodevelopmental disorders than their peers without CHD, and at 22 months of age score significantly lower on measures of cognitive, motor and language development than the standardised population mean [5]. Additionally, children and young adults with CHD face an elevated risk of developing cancer [6]. Hence, CHD not only impairs cardiovascular function but also causes co-morbidities that profoundly affect quality of life and life expectancy.

Due to the complexity in genetic and cellular interactions needed for the heart to form properly, perhaps it is not surprising that many cases of CHD result from pathological genetic variation. However, despite much research into the underlying genetic associations with CHD, a specific genetic diagnosis is still lacking in up to 80% of sporadic CHD cases and up to 70% of familial CHD cases [7, 8]. Approximately two-thirds of CHD cases occur due to complex genetic mechanisms that involve interactions between multiple genes [9], adding challenges to the process of confirming pathogenicity. Moreover, CHD can arise from *de novo* mutations, with the absence of familial inheritance [10]. Previous studies have estimated that several hundred genes may contribute to CHD susceptibility [11]. Genome sequencing of patients with CHD has identified genetic variants in genes not previously known to play a part in cardiac development [11–13]. However, a standardised system to evaluate the significance of these variants is lacking, despite the urgent need for such knowledge to facilitate genetic diagnosis and inform genetic counselling.

Advancements in discovering further genetic causes for CHDs can be limited by the challenges in prioritising candidate genes from CHD patient sequence analyses. Comparison to mutant mouse models can aid in prioritisation, however data from mouse knockout experiments are limited. As of April 2024, the International Mouse Phenotyping Consortium (IMPC) [14] completed phenotyping for knockouts of only 8707 of the approximately 22,500 protein-coding mouse genes, leaving many genes uncharacterised. Consequently, current sequence variant prioritisation methods often fail due to a lack of data, lack of knowledge over which factors are most informative for determining pathogenicity, and challenges in manually integrating information from disparate data sources [15, 16]. A new method to prioritising candidate genes is imperative, and machine learning (ML), a subset of artificial intelligence (AI), holds the potential to address this challenge.

Machine learning uses algorithms to enable computers to find patterns in data characteristic of a group and to make predictions based on those patterns. Since machine learning approaches have not yet been applied to CHD gene prediction in mammalian species, we developed a supervised machine learning classifier to identify characteristics of genes required for cardiac development in mice. These genes, when mutated, could serve as promising candidate disease genes for CHDs in humans. Utilising a training dataset comprising genes known to be either indispensable for cardiac development or not required, we trained a Random Forest classifier to distinguish between these two classes. The efficacy of our method was validated by classifying genes independent of the training dataset. We then predicted the cardiac development association status of all protein-coding genes in the mouse genome. Large-scale bioinformatic analyses revealed functions linking the predicted cardiac genes to cardiac development. RNA-Seq data showed similar expression patterns between our predicted and known cardiac genes. Many of our predicted genes are orthologues of newly published human CHD genes, affirming that our classifier predictions do indeed include human CHD gene candidates. The knowledge of cardiac development genes may assist gene prioritisation from sequence analysis, potentially accelerating the diagnosis of CHDs.

## Results

### Datasets

The Mouse Genome Informatics (MGI) database [17] serves as a comprehensive repository for published gene data on mouse knockout phenotypes. Initially, we compiled two datasets from MGI: one containing 1415 mouse genes known to be involved in cardiac development (cardiac genes), and another containing 6808 genes known not to be involved in cardiac development (non-cardiac genes). These classifications were based on phenotype annotations derived from null alleles of targeted single-gene knockouts. To ensure precise gene classification, genes ambiguously labelled in MGI as both cardiac and non-cardiac were manually reviewed against published literature to verify their roles in cardiac development. To focus on features specific to protein function, we limited our analysis to protein-coding genes. As e result, after excluding 173 cardiac and 235 non-cardiac genes, our final datasets comprised 1242 cardiac genes and 6573 non-cardiac genes (Tables S1 and S2).

### Cardiac Genes are Different in Features Compared to Non-cardiac Genes

We gathered a wide range of features of mouse protein-coding genes, encompassing gene and protein sequence-based attributes, gene expression, gene ontology (GO) annotations, and protein-protein interaction details. In total, we analysed 127 features (detailed in the Supporting Information) to compare cardiac and non-cardiac genes and identify properties associated with cardiac development. This analysis revealed significant differences in the values of several features between the cardiac and non-cardiac datasets (Table 1).

**Table 1.**
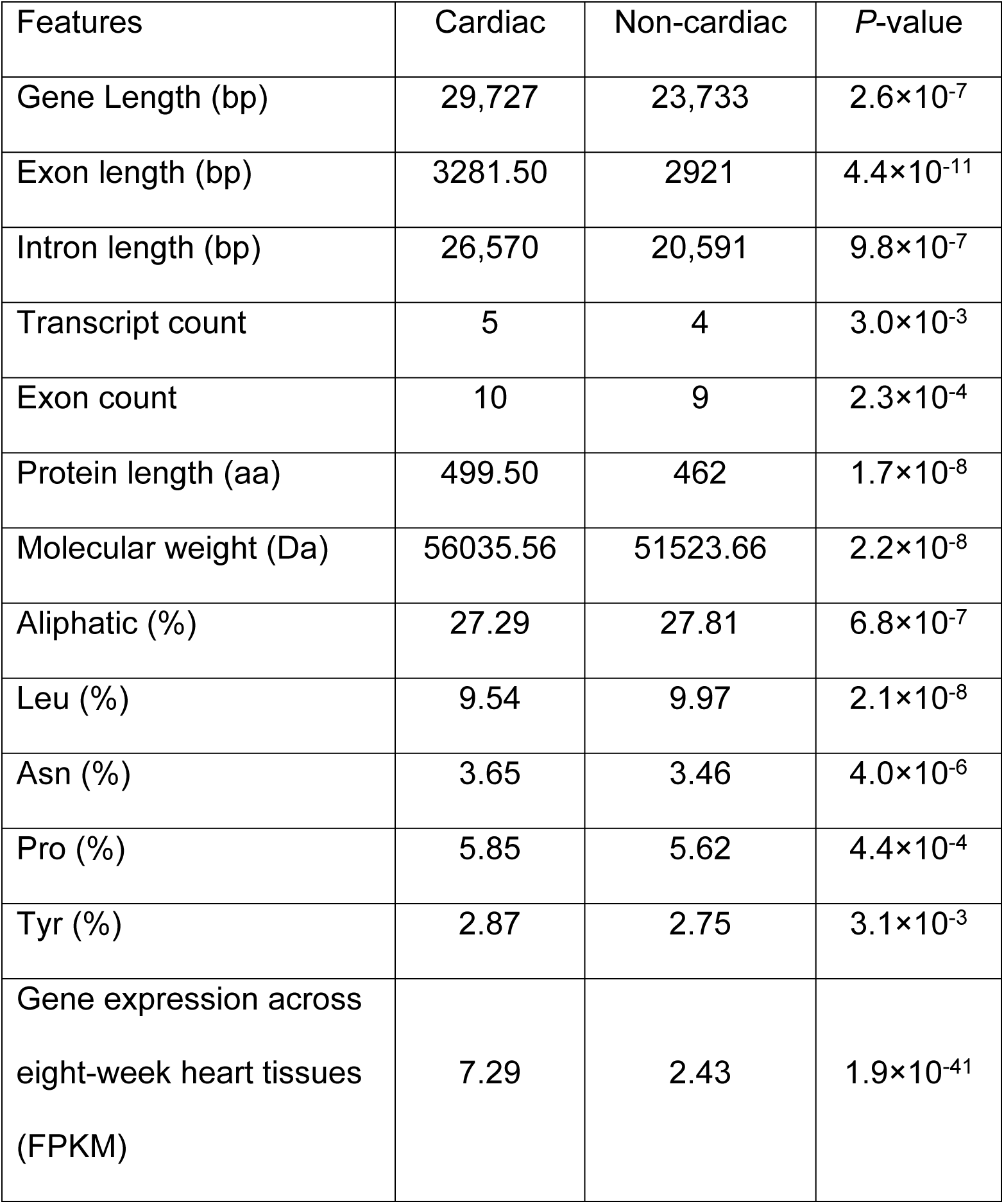
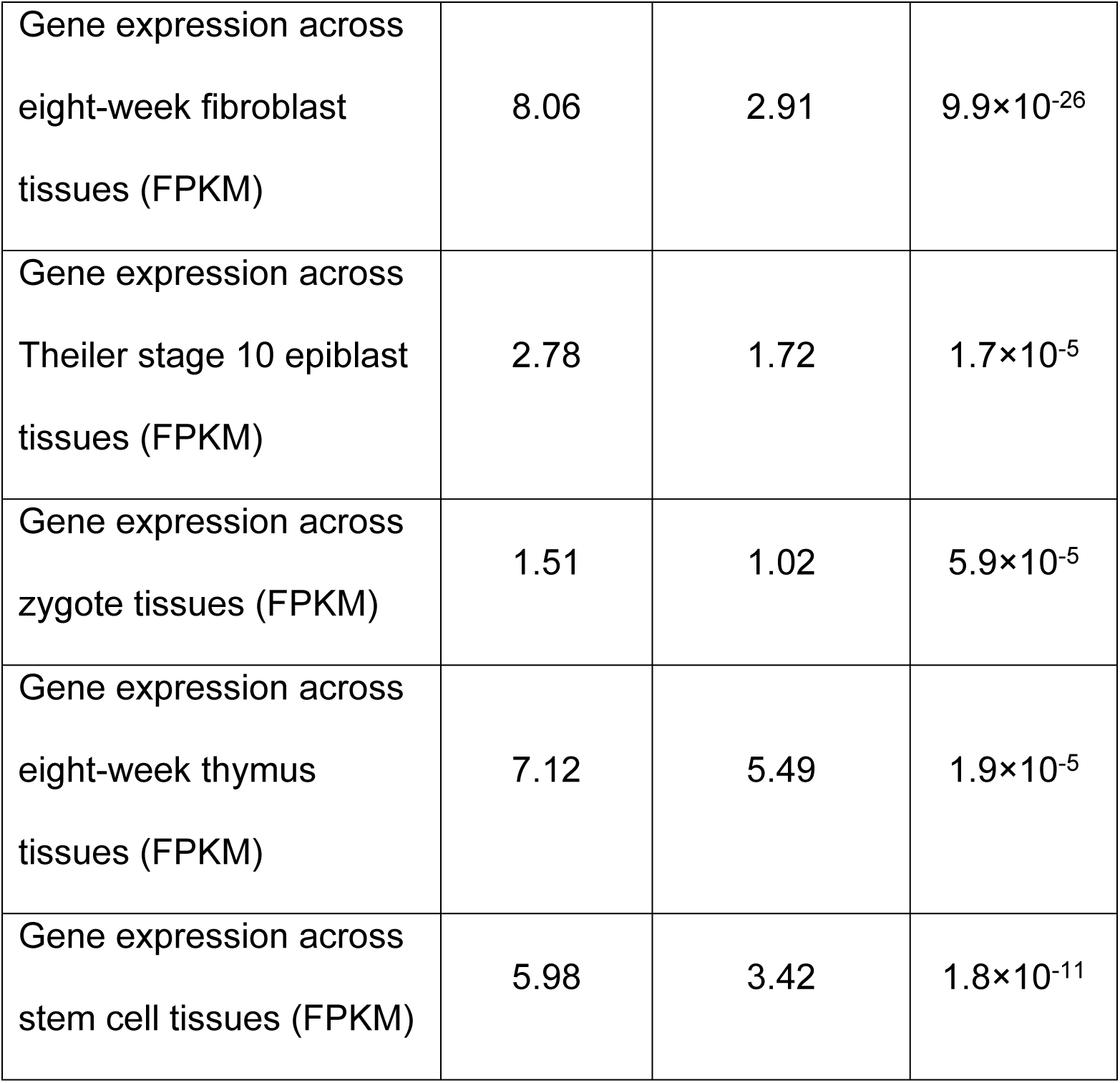
List of statistically significant features between cardiac and non-cardiac genes. The median value of each feature is reported. *P*-values were determined from the Mann-Whitney U test. Statistically significant results were evaluated based on the Bonferroni corrected *P*-value.

| Features | Cardiac | Non-cardiac | *P*-value |
| --- | --- | --- | --- |
| Gene Length (bp) | 29,727 | 23,733 | $2.6 \times 10^{-7}$ |
| Exon length (bp) | 3281.50 | 2921 | $4.4 \times 10^{-11}$ |
| Intron length (bp) | 26,570 | 20,591 | $9.8 \times 10^{-7}$ |
| Transcript count | 5 | 4 | $3.0 \times 10^{-3}$ |
| Exon count | 10 | 9 | $2.3 \times 10^{-4}$ |
| Protein length (aa) | 499.50 | 462 | $1.7 \times 10^{-8}$ |
| Molecular weight (Da) | 56035.56 | 51523.66 | $2.2 \times 10^{-8}$ |
| Aliphatic (%) | 27.29 | 27.81 | $6.8 \times 10^{-7}$ |
| Leu (%) | 9.54 | 9.97 | $2.1 \times 10^{-8}$ |
| Asn (%) | 3.65 | 3.46 | $4.0 \times 10^{-6}$ |
| Pro (%) | 5.85 | 5.62 | $4.4 \times 10^{-4}$ |
| Tyr (%) | 2.87 | 2.75 | $3.1 \times 10^{-3}$ |
| Gene expression across eight-week heart tissues (FPKM) | 7.29 | 2.43 | $1.9 \times 10^{-41}$ |
| Gene expression across eight-week fibroblast tissues (FPKM) | 8.06 | 2.91 | $9.9 \times 10^{-26}$ |
| Gene expression across Theiler stage 10 epiblast tissues (FPKM) | 2.78 | 1.72 | $1.7 \times 10^{-5}$ |
| Gene expression across zygote tissues (FPKM) | 1.51 | 1.02 | $5.9 \times 10^{-5}$ |
| Gene expression across eight-week thymus tissues (FPKM) | 7.12 | 5.49 | $1.9 \times 10^{-5}$ |
| Gene expression across stem cell tissues (FPKM) | 5.98 | 3.42 | $1.8 \times 10^{-11}$ |

We observed that cardiac genes exhibit distinct characteristics compared to non-cardiac genes, as depicted in Table 1 and Fig 1. Cardiac genes are more likely to be long, have multiple transcripts, a greater number of exons, and possess longer exons and introns. A higher proportion of cardiac genes are expressed during critical stages of mouse development, including blastocyst (44.7% vs. 39.5%, Chi-squared *P-*value = 8.2 × 10^-3^), gastrula (42.4% vs. 37.4%, Chi-squared *P-*value = 8.2 × 10^-3^), organogenesis (72.1% vs. 61.3%, Chi-squared *P-*value = 1.1 × 10^-5^), and neonate (80.8% vs. 73.1%, Chi-squared *P-*value = 3.5 × 10^-3^) stages, compared to non-cardiac genes. Additionally, cardiac genes demonstrate higher expression levels in various developmental tissues, including eight-week heart, eight-week fibroblast, Theiler stage 10 epiblast, zygote, eight-week thymus, and stem cell tissues (Table 1).

**Fig 1:**
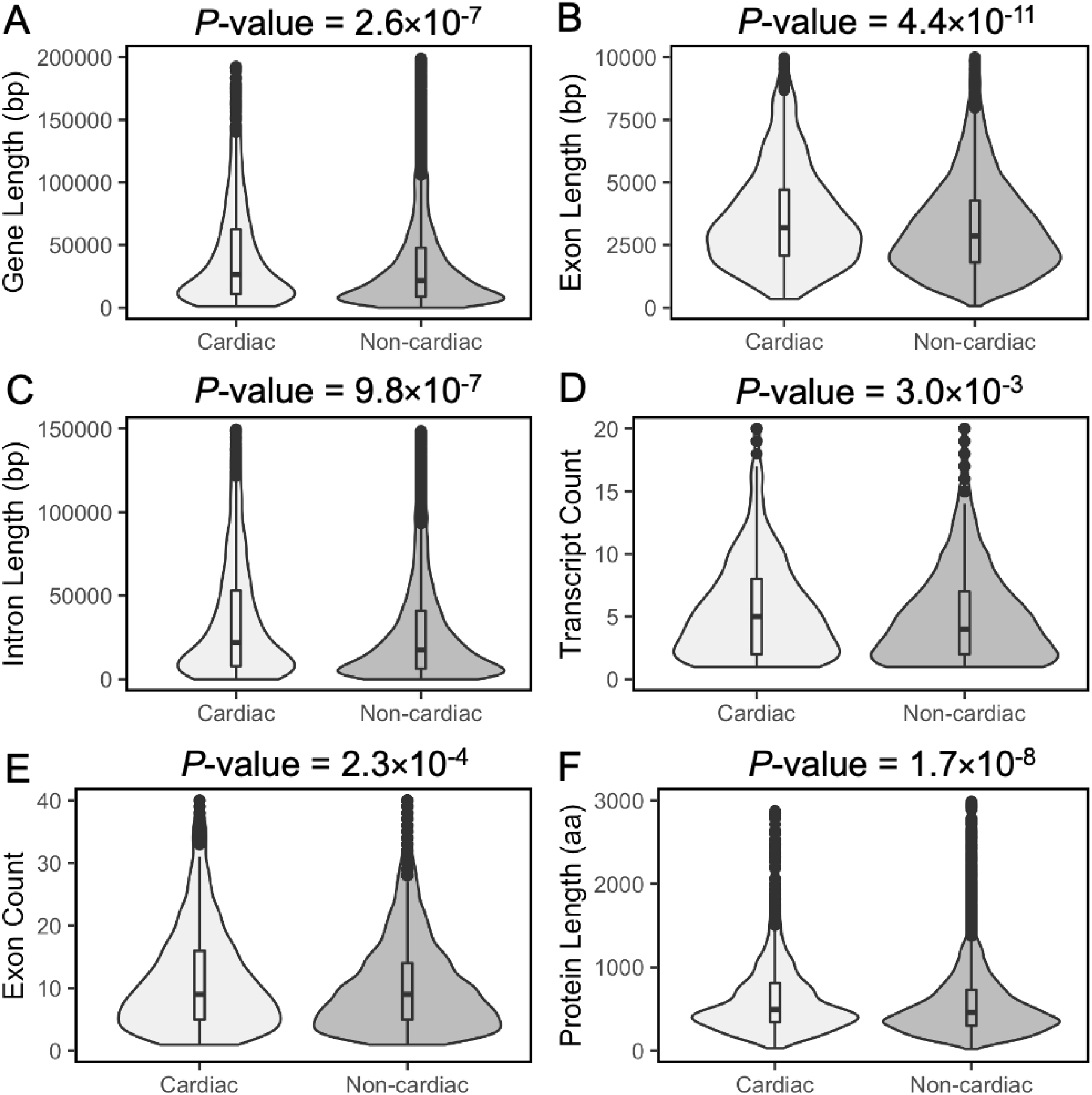
Violin plots showing the distributions of total gene length, exon length, intron length, transcript count, exon count and protein length in cardiac and non-cardiac datasets. Horizontal lines within boxplots represent median values. *P*-values were determined using the Mann-Whitney *U* test and adjusted for multiple comparisons with the Bonferroni correction.

Moreover, cardiac genes are characterised by having low probability of loss-of-function (pLoF) scores and high probability of being loss-of-function intolerant (pLI) in gnomAD, as illustrated in Fig 2. The gene pLoF score is a metric used to assess the likelihood of observing variants resulting in loss-of-function of a particular gene, based on the data collected in gnomAD v2.1.1. Therefore, a low pLoF score means a gene is less likely to have loss-of-function variants present in sequence data compiled in gnomAD. On the other hand, the gene pLI score is used to measure the tolerance of a gene to loss-of-function mutations. It quantifies the likelihood that a gene is essential for normal development and survival. A high pLI score indicates that a gene is likely intolerant to loss-of-function mutations and may have critical functions. A gene with a high pLI will not tolerate loss-of-function (LoF) variants; hence, it will have a low pLoF.

**Fig 2:**
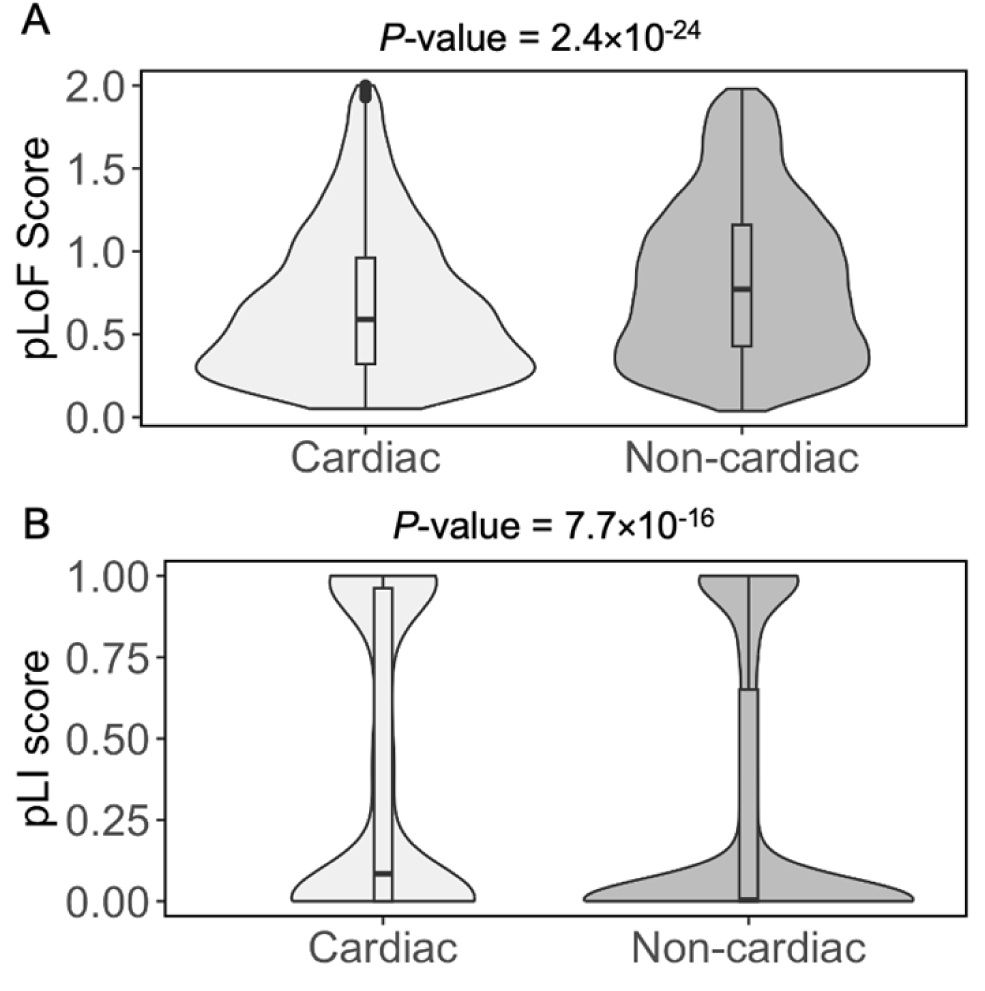
Violin plots showing the distributions of pLoF and pLI scores in cardiac and non-cardiac datasets. Horizontal lines within boxplots represent median values. *P*-values were determined using the Mann-Whitney U test and evaluated for significance with Bonferroni correction.

Analyses of mouse protein sequence data revealed that cardiac genes are more likely to encode proteins with longer lengths (Fig 1f) and higher molecular weights (Table 1). These proteins exhibit higher proportions of Asparagine (Asn), Proline (Pro), and Tyrosine (Tyr) residues, while non-cardiac proteins display higher proportions of aliphatic and Leucine (Leu) residues (Table 1). Moreover, our analysis revealed that cardiac proteins exhibit enrichment in specific functional annotations compared to non-cardiac proteins. Cardiac proteins are more likely to function as oxidoreductases (4.8% vs. 3.2%, Chi-squared *P-*value = 7.9 × 10^-3^), transcription factors (19.1% vs. 11.9%, Chi-squared *P-*value = 1.9 × 10^-10^), phosphorylated proteins (58.2% vs. 46.3%, Chi-squared *P-*value = 3.2 × 10^-8^) and acetylated proteins (20.6% vs. 15.7%, Chi-squared *P-*value = 1.1 × 10^-4^), and have a higher frequency of signal peptide motifs (28.1% vs. 22.4%, Chi-squared *P-*value = 1.3 × 10^-4^). Furthermore, subcellular localisation analysis utilising UniProt annotation revealed that a greater percentage of cardiac proteins are localised within the nucleus (33.1%. vs 27.1%, Chi-squared *P-*value = 2.6 × 10^-4^) and extracellular region (16.4% vs 10.9%, Chi-squared *P-*value = 2.5 × 10^-7^) compared to non-cardiac proteins, which are mostly localised in the cytoplasm and membrane.

There is an emerging view that additional genes associated with biological functions and phenotypes can be identified due to “guilt by association” with genes already known to participate in those processes [18–23]. Most approaches investigating gene associations have used protein interaction data [19–23], although others have used similarity of functional annotation [18]. In our study, we investigated the protein-protein interaction (PPI) data for mouse proteins from the I2D [24] database to determine whether the cardiac PPI network exhibits distinct network properties compared to the non-cardiac network. We focused solely on known mouse PPIs to ensure the inclusion of high-quality interactions. For each cardiac and non-cardiac protein, we computed 13 network properties to assess their significance within their respective PPI networks. Cardiac proteins demonstrated a higher number of interactions (higher degrees) compared to non-cardiac proteins within their interaction network (Fig 3A). Moreover, the betweenness centrality, serving as an indicator of a protein’s centrality within the PPI network, was significantly higher for cardiac proteins than non-cardiac proteins (Fig 3B). Additionally, we observed significantly higher closeness centrality values for cardiac proteins compared to non-cardiac proteins (Fig 3C).

**Fig 3:**
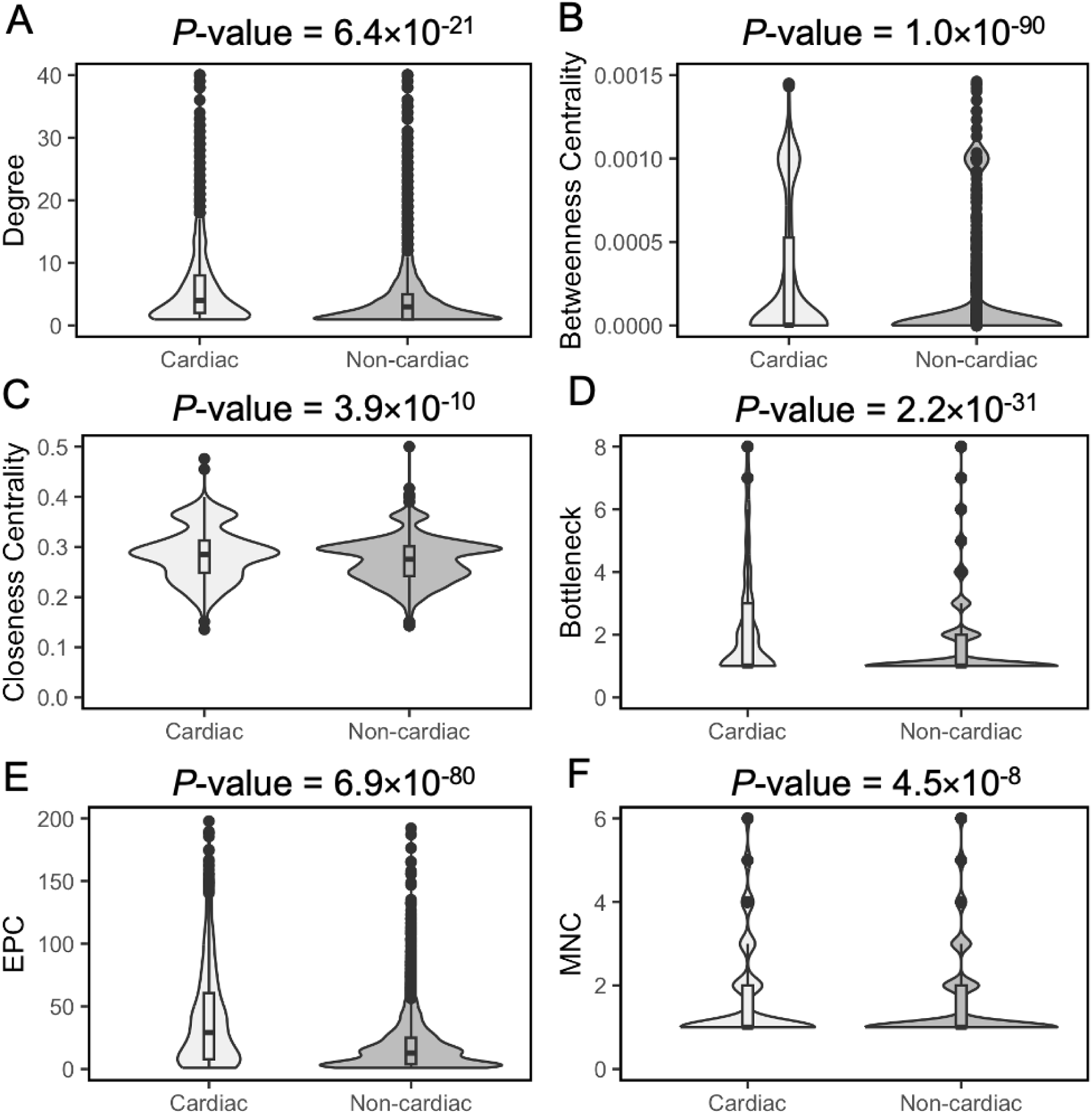
Violin plots showing the distributions of **PPI** network features **of** cardiac and non-cardiac proteins. Degree (A), Betweennep-ss centrality (B), Closeness centrality (C), Bottleneck (D), Edge Percolation Component (E) and Maximum Neighbourhood Component (F) of cardiac and non-cardiac proteins in the PPI networks. Horizontal lines within boxplots represent median values. *P*-values were determined using the Mann-Whitney U test and evaluated for significance with Bonferroni correction.

To identify protein nodes with large number of interactions (hubs) in the PPI network, we utilised the Hub object Analyzer (Hubba) [25]. This tool allowed us to explore four additional network properties: BottleNeck (BN), Edge Percolation Component (EPC), Maximum Neighbourhood Component (MNC), and Density of Maximum Neighbourhood Component (DMNC). These properties define probable hubs, or highly connected proteins, in the interaction network. Our investigation revealed that cardiac proteins tend to have high BN, EPC and MNC values compared to non-cardiac proteins, as illustrated in Fig 3.

We analysed the Gene Ontology (GO) terms [26] associated with genes in our datasets using the web-based tool DAVID [27], as GO is the most widely used approach for annotating gene functions. Our analysis revealed notable differences in the GO term annotations for biological process and cellular component classes between cardiac and non-cardiac gene groups. For biological processes, GO data analysis demonstrated that cardiac genes are primarily involved in processes related to ‘heart development’, ‘heart morphogenesis’, ‘angiogenesis’, ‘vasculogenesis’, ‘heart looping’ and in utero embryonic development’ (Table 2). In contrast, non-cardiac genes are predominantly associated with processes like ‘immune system response’, ‘ion transport’, ‘cell adhesion’, ‘inflammatory response’, ‘cell adhesion’ and ‘brain development’ (Table 3). This result also validates the selection of our cardiac gene dataset. For cellular component, terms most frequently associated with cardiac genes include ‘cell surface’, ‘cytoplasm’, ‘extracellular region’, ‘plasma membrane” and ‘nucleus’’. On the other hand, the non-cardiac dataset exhibited enrichment for terms such as ‘glutamatergic synapse’, ‘synapse’, ‘cell junction’, ‘neuron projection’, and ‘cytosol’. Detailed lists of the 10 most enriched GO terms for cellular components are provided in Tables S3–S4.

**Table 2:** Top 10 GO terms for mouse cardiac development genes related to biological processes, identified using the DAVID Functional Annotation tool.

| Term ID | Term | Count | % | *P*-value | Bonferroni |
| --- | --- | --- | --- | --- | --- |
| GO:0007507 | heart development | 164 | 13.16 | $1.34 \times 10^{-116}$ | $9.39 \times 10^{-113}$ |
| GO:0001525 | angiogenesis | 123 | 9.87 | $2.59 \times 10^{-72}$ | $1.82 \times 10^{-68}$ |
| GO:0001570 | vasculogenesis | 52 | 4.17 | $1.20 \times 10^{-43}$ | $8.40 \times 10^{-40}$ |
| GO:0001568 | blood vessel development | 51 | 4.09 | $2.78 \times 10^{-41}$ | $1.95 \times 10^{-37}$ |
| GO:0001701 | in utero embryonic development | 99 | 7.95 | $4.93 \times 10^{-39}$ | $3.46 \times 10^{-35}$ |
| GO:0007275 | multicellular organism development | 188 | 15.09 | $9.96 \times 10^{-36}$ | $7.00 \times 10^{-32}$ |
| GO:0003007 | heart morphogenesis | 45 | 3.61 | $8.91 \times 10^{-35}$ | $6.26 \times 10^{-31}$ |
| GO:0003151 | outflow tract morphogenesis | 38 | 3.05 | $7.44 \times 10^{-31}$ | $5.22 \times 10^{-27}$ |
| GO:0006468 | protein phosphorylation | 118 | 9.47 | $6.25 \times 10^{-29}$ | $4.39 \times 10^{-25}$ |
| GO:0001947 | heart looping | 39 | 3.13 | $3.58 \times 10^{-28}$ | $2.51 \times 10^{-24}$ |

**Table 3:** Top 10 GO terms for mouse non-cardiac genes related to biological processes, identified using the DAVID Functional Annotation tool.

| Term ID | Term | Count | % | *P*-value | Bonferroni |
| --- | --- | --- | --- | --- | --- |
| GO:0002376 | immune system process | 299 | 4.54 | $7.40 \times 10^{-38}$ | $7.27 \times 10^{-34}$ |
| GO:0030154 | cell differentiation | 498 | 7.56 | $1.62 \times 10^{-33}$ | $1.60 \times 10^{-29}$ |
| GO:0007165 | signal<br>transduction | 563 | 8.55 | $1.27 \times 10^{-27}$ | $1.25 \times 10^{-23}$ |
| GO:0006811 | ion transport | 310 | 4.71 | $7.15 \times 10^{-25}$ | $7.03 \times 10^{-21}$ |
| GO:0007399 | nervous system<br>development | 242 | 3.67 | $1.23 \times 10^{-24}$ | $1.21 \times 10^{-20}$ |
| GO:0007268 | chemical<br>synaptic<br>transmission | 129 | 1.96 | $9.73 \times 10^{-24}$ | $9.57 \times 10^{-20}$ |
| GO:0006954 | inflammatory<br>response | 213 | 3.23 | $7.30 \times 10^{-23}$ | $7.18 \times 10^{-19}$ |
| GO:0007275 | multicellular<br>organism<br>development | 512 | 7.77 | $2.53 \times 10^{-22}$ | $2.49 \times 10^{-18}$ |
| GO:0007283 | spermatogenesis | 260 | 3.95 | $6.01 \times 10^{-21}$ | $5.91 \times 10^{-17}$ |
| GO:0007218 | neuropeptide<br>signaling<br>pathway | 67 | 1.02 | $2.15 \times 10^{-16}$ | $2.18 \times 10^{-12}$ |

### Processing Datasets for Machine Learning

We identified distinctive features that could effectively discriminate between cardiac and non-cardiac genes. We therefore sought to develop a machine learning classifier that could categorize a mouse gene as either cardiac or non-cardiac based on these features (Fig 4). Using 127 features as input, we generated training datasets for classification. The quality of the training dataset is crucial for the success of machine learning classifiers. Initially, our dataset included 1242 cardiac genes and 6573 non-cardiac mouse genes, resulting in a significantly imbalanced class frequency ratio of 1:5.3. To mitigate the impact of class imbalance in the training dataset, we created balanced training datasets. These included all 1242 cardiac genes and an equal number of randomly selected non-cardiac genes (1242 out of 6573 total non-cardiac genes). This approach ensured our classifier was effectively trained without bias towards the majority class [28, 29].

**Fig 4:**
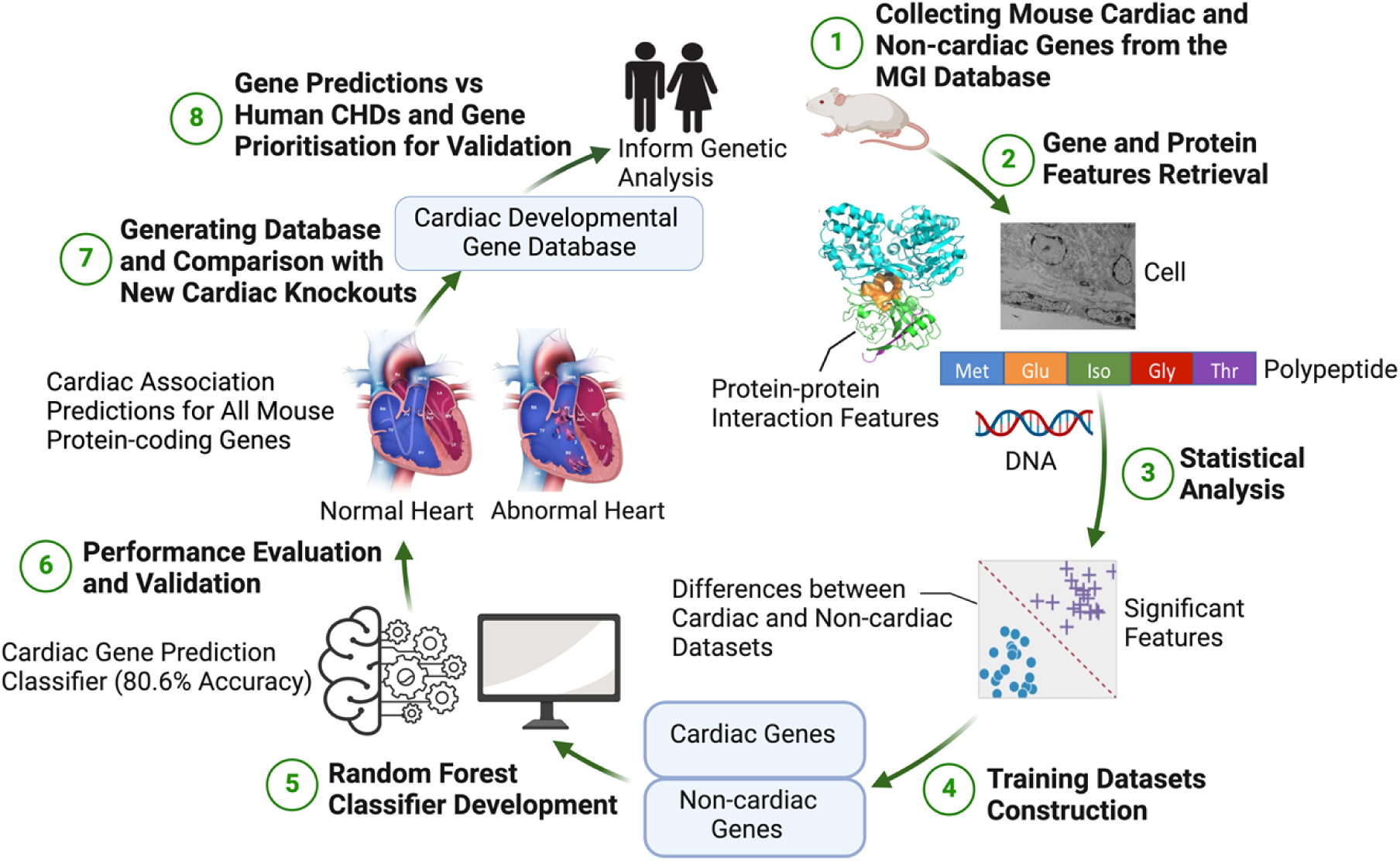
The workflow for predicting mouse cardiac development genes from gene and protein sequence features using a Random Forest classification model. First, features of mouse genes are collected from publicly available databases. Subsequently, statistical analyses and feature selection were performed to pinpoint the most informative features distinguishing between known cardiac and non-cardiac genes. A Random Forest classifier was constructed to predict cardiac and non-cardiac genes. Finally, this classifier was deployed to predict cardiac development association status for all protein coding genes within the mouse genome which were not used in developing the classifier. A database of prediction results has been developed and is available to the public.

We also constructed test datasets comprising mouse genes not included in the training datasets to assess the accuracy of our machine learning classifier. Two test datasets were compiled for the validation of our predictions. Test dataset 1 (Table S5) consists of 5331 mouse knockout genes selected randomly from our non-cardiac dataset, which were not included in our initial training set selection. Test dataset 2 (Table S6) contains 1193 mouse reporter line mutants. We restricted our training datasets to genes with mouse targeted deletion alleles only, so genes with reporter mutant lines were excluded from the compilation of our training datasets. However, phenotypic data are available in MGI for these reporter lines, and therefore they can be used as a second test set of genes with experimentally known cardiac or non-cardiac development status.

### Performance of the Random Forest Classifier

To develop our classifier, we employed the Random Forest (RF) [30] method implemented using open source machine learning software in WEKA [31], to construct a machine learning classifier aimed at identifying genes likely to be involved in cardiac development. RF, an ensemble classifier composed of multiple decision tree models, has been demonstrated to exhibit high accuracy across various studies [32–36]. Our initial RF classifier (RF-1) was trained on all features of the training dataset using a 10-fold cross-validation approach. This methodology was employed to enhance the robustness of our classifier and mitigate potential overfitting issues. The cross-validation accuracy of RF-1 classifier was 79.8% (1982/2484), with 951 true-positives (TPs), 291 false-negatives (FNs), 1031 true-negatives (TNs), and 211 false-positives (FPs). The performance of this classifier was also assessed using various performance measures, the detailed values of which are presented in Table 4 for each class.

**Table 4:** 10-fold cross validation performance of the Random Forest classifiers, trained and evaluated on the training dataset. Data from before and after feature selection are presented. Here, TP = True Positive; FP = False Positive; ROC = Receiver Operating Curve; PRC = Precision-Recall Curve.

| Classifiers | Accuracy (%) | Gene Class | TP Rate | FP Rate | Precision | F-Measure | ROC area (AUC) | PRC area |
| --- | --- | --- | --- | --- | --- | --- | --- | --- |
| RF-1 | 79.8 | Cardiac | 0.766 | 0.170 | 0.818 | 0.791 | 0.870 | 0.866 |
|  |  | Non-cardiac | 0.830 | 0.234 | 0.780 | 0.804 | 0.870 | 0.867 |
|  |  | Weighted average | 0.798 | 0.202 | 0.799 | 0.798 | 0.870 | 0.866 |
| RF-2 | 80.6 | Cardiac | 0.791 | 0.180 | 0.815 | 0.803 | 0.882 | 0.881 |
|  |  | Non-cardiac | 0.820 | 0.209 | 0.797 | 0.809 | 0.882 | 0.879 |
|  |  | Weighted average | 0.806 | 0.194 | 0.806 | 0.806 | 0.882 | 0.880 |

Using all features in a training dataset is not always optimal for classifier performance. Feature selection can reduce overfitting, enhance classification accuracy, and speed up training. We employed the Information Gain feature selection method in WEKA to identify the most relevant mouse gene features for classification, ultimately selecting 73 features from the original set of 127 (Table 5 and S7). This process resulted in the selection of a subset of 73 features from the original set of 127 (Table 5 and S7). Most of these selected features exhibited statistically significant differences in values between cardiac and non-cardiac genes, confirming their discriminative potential in our study. We developed another RF classifier (RF-2) based on the 73 selected features, employing 10-fold cross-validation. The accuracy of this classifier improved to 80.59%, with 983 TPs, 259 FNs, 1019 TNs, and 223 FPs. Performance metrics presented in Table 4 confirm that feature selection significantly enhanced RF classifier’s performance.

**Table 5:** Top 10 features selected from the training dataset using the Information Gain feature selection method. Features are sorted in descending order with respect to the corresponding information gain value, with the most informative feature listed first. Many of the informative features are associated with protein interaction networks.

| Information Gain | Feature | Feature Type | Observation for Cardiac Genes |
| --- | --- | --- | --- |
| 0.09884 | Eigenvector | PPI Network | Low |
| 0.09187 | Betweenness Centrality | PPI Network | High |
| 0.08637 | Edge Percolation Component | PPI Network | High |
| 0.0851 | Average Shortest Path Length | PPI Network | Low |
| 0.08106 | Closeness Centrality | PPI Network | High |
| 0.04575 | Degree | PPI Network | High |
| 0.04199 | Bottleneck | PPI Network | High |
| 0.03697 | Clustering Coefficient | PPI Network | High |
| 0.03679 | GO:0007507<br>Heart Development | Biological<br>Process | High proportion |
| 0.02889 | Eight-week Heart Expression | Developmental<br>Expression<br>Pattern | Expressed in<br>high proportion |
| 0.0208 | pLoF | Gene<br>Essentiality | Low |

To assess potential overfitting of the Random Forest classifier due to the training dataset, we generated four additional balanced training datasets, each containing different subsets of non-cardiac genes. We then trained a separate Random Forest classifier on each of these datasets. The performance evaluations, detailed in Table S8, revealed a mean accuracy of 80.1% with a standard deviation of 0.6. These results confirm that the Random Forest classifier is not biased by the choice of the training dataset.

The performance of our classifier was then evaluated using two test datasets. Test dataset 1 contained all non-cardiac genes with known experimental data that had not been included in training the RF classifier; 86% of the genes (4603 out of a total of 5331 genes) were correctly identified as non-cardiac in this dataset. Moreover, our classifier showed a very strong performance on Test dataset 2, correctly identifying the cardiac development status for 88% of the genes (1052 out of 1193 genes total) with phenotypic data from mouse Cre reporter line experiments.

### Cardiac Development Associated Gene Prediction

We applied our RF classifier to the full list of 12,375 mouse protein-coding genes (retrieved from the MouseMine [37] database) with unknown cardiac-association status (Table S9) and generated cardiac development status predictions for all these genes. We identified 4472 (36% of the prediction dataset) mouse genes likely to play a role in cardiac development. A total of 7901 (64% of the prediction dataset) genes are predicted not to be associated with cardiac development. We ranked these genes by the confidence level of their prediction (Table S10). The top 15 predicted cardiac genes are listed in Table 6. The expression of the top 15 predicted cardiac genes was confirmed by RT-PCR analysis using RNA from E10.5 mouse whole embryos, E15.5 mouse embryonic hearts, and adult mouse hearts (Figure S1).

**Table 6:** Top 15 mouse cardiac development genes predicted by our Random Forest classifier.

| **Gene Name** | **Encoded protein and function** | **Predicted Class** | **Confidence** |
| --- | --- | --- | --- |
| Epn2 | Epsin-2; involves in clathrin-coated endocytosis | Cardiac | 0.995 |
| Gne | Glucosamine (UDP-N-acetyl)-2-epimerase; initiates the biosynthesis of N-acetylneuraminic acid (NeuAc), a key precursor for sialic acids | Cardiac | 0.995 |
| Sp3 | Transcription factor Sp3; act as an activator or repressor depending on isoform or modifications | Cardiac | 0.99 |
| Cdk2 | Cyclin-dependent kinase 2; involves in the control of the cell cycle | Cardiac | 0.98 |
| Bmi1 | Polycomb complex protein BMI-1; requires maintaining the transcriptionally repressive state of many genes | Cardiac | 0.98 |
| Ccne1 | G1/S-specific cyclin-E1; Essential for the control of the cell cycle at the G1/S (start) transition | Cardiac | 0.975 |
| Dvl1 | Segment polarity protein dishevelled homolog DVL-1; participates in Wnt signaling | Cardiac | 0.975 |
| Qki | KH domain-containing RNA-binding protein QKI; regulates RNA metabolic processes | Cardiac | 0.975 |
| Dpysl2 | Dihydropyrimidinase-related protein 2; plays a role in neuronal development and polarity | Cardiac | 0.97 |
| Cenpc1 | Centromere protein C; plays a central role in assembly of kinetochore proteins | Cardiac | 0.97 |
| Gna12 | Guanine nucleotide-binding protein subunit alpha-12; involves as modulators or transducers in various transmembrane signaling systems | Cardiac | 0.97 |
| Dnmt1 | DNA (cytosine-5)-methyltransferase 1; essential for epigenetic inheritance | Cardiac | 0.965 |
| Atf7ip | Activating transcription factor<br>7-interacting protein 1;<br>modulates transcription<br>regulation and chromatin<br>formation | Cardiac | 0.965 |
| Vps26a | Vacuolar protein sorting-<br>associated protein 26A; acts<br>as a component of the<br>retromer cargo-selective<br>complex (CSC) | Cardiac | 0.965 |
| Enah | ENAH actin regulator; involves<br>in cytoskeleton remodeling and<br>cell polarity | Cardiac | 0.965 |

Consistent with their cardiac development prediction status, all the top 15 predicted genes exhibited expression in these tissues.

### Cardiac Development Gene Database

We developed a publicly accessible database, CDGD (Cardiac Development Gene Database; URL: http://130.88.96.175/), which contains information regarding the cardiac/non-cardiac status of all protein-coding genes within the mouse genome, derived either from published literature or our predictions. The database contains confidence scores indicating the predicted probabilities of the genes to be cardiac or non-cardiac. The database can be searched for known or predicted mouse genes using various identifiers such as gene name, MGI ID, Ensembl ID, and UniProt ID. Furthermore, users can retrieve lists of all cardiac and non-cardiac genes (both known and predicted) within the mouse genome or within specific chromosomal and genomic regions. All search results are downloadable as CSV files for further analysis.

### PPI Networks of Cardiac and Non-cardiac Genes

Since we found protein interaction network features to be highly informative for our classifier development, we decided to compare the topology features of protein interaction networks involving proteins whose cardiac status was known and those networks involving proteins with predicted status. We utilised PPI databases BioGRID [38] and STRING [39] along with I2D to extract protein interaction data for the encoded proteins of both predicted and known genes. Invoking the ‘guilt by association’ principle, STRING data has been previously utilised to successfully predict novel CHD genes based on interactions with known CHD proteins [13], therefore, inclusion of interaction data from STRING is highly relevant to our cardiac gene predictions analysis. Based on the guilt-by-association principle, we would expect a greater number of interactions between known and predicted cardiac proteins. Our observations confirmed that known cardiac proteins exhibit more interactions with predicted cardiac proteins than with predicted non-cardiac proteins. This trend was consistently observed across all three protein-protein interaction (PPI) databases (Table S11). Furthermore, we constructed four distinct PPI networks from each database: K-Cardiac (encompassing PPIs for all known cardiac proteins), K-NC (encompassing PPIs for all known non-cardiac proteins), P-Cardiac (encompassing PPIs for all predicted cardiac proteins), and P-NC (encompassing PPIs for all predicted non-cardiac proteins) networks. Using the ‘network analyzer’ plugin of Cytoscape v3.9.1, we computed the network properties for these networks. Our analysis revealed that known cardiac proteins exhibited a higher number of interactions (higher degrees) compared to known non-cardiac proteins within their respective interaction networks, consistent across both BioGRID and STRING (Fig 5), and mirroring observations made with I2D. Furthermore, we found a similar trend with the PPI networks generated from our predicted cardiac proteins, which also displayed higher degrees compared to networks of predicted non-cardiac proteins. These trends remained consistent across all PPI databases, as confirmed by Mann-Whitney U statistical tests (Fig 5). Notably, many of the predicted cardiac proteins have fewer interactions compared to known cardiac proteins, yet they were still predicted to be associated with cardiac development. One reason for this could be that they are less studied.

**Fig 5:**
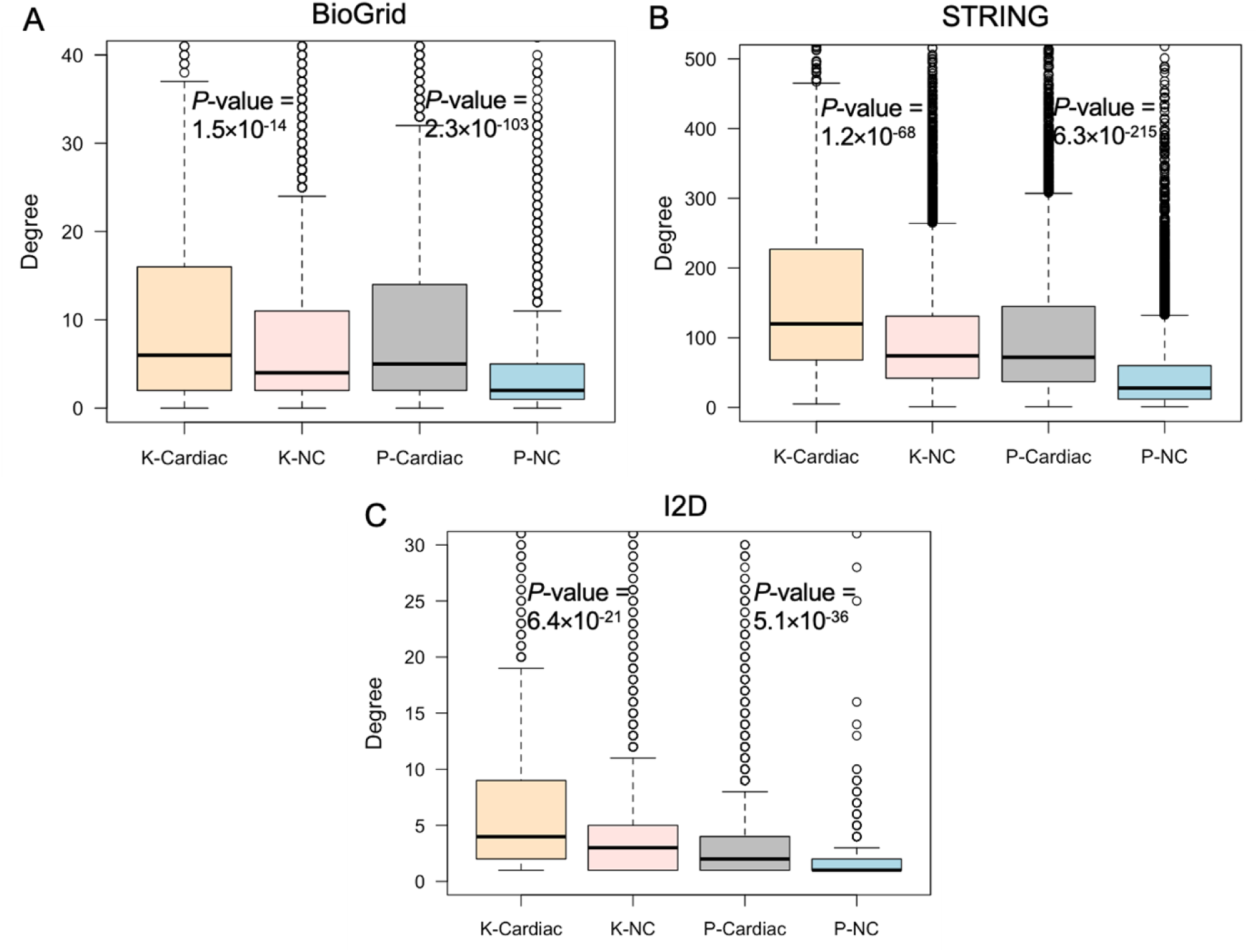
Degree of cardiac and non-cardiac proteins within the PPI networks sourced from BioGRID (A), STRING (B) and I2D (C) databases. Here, K-Cardiac represents the PPI network of known cardiac proteins; K-NC represents the PPI network of known non-cardiac proteins; P-Cardiac represents the PPI network of predicted cardiac proteins; and P-NC represents the PPI network of predicted non-cardiac proteins. *P*-values were determined using the Mann-Whitney U test.

### Developmental Expression Patterns of Cardiac and Non-cardiac Genes

Genes that are involved in the same biological process would also be expected to have similarities in expression – in concordance with ‘guilt by association’ [40]. Therefore, we hypothesised that genes in our prediction datasets should show similarities in developmental expression pattern (co-expression) to genes in our known datasets for each gene class. To test this hypothesis, we analysed developmental gene expression profiles across four distinct categories of gene pairs: (1) predicted cardiac vs known cardiac, (2) predicted cardiac vs known non-cardiac, (3) predicted non-cardiac vs known cardiac, and (4) predicted non-cardiac vs known non-cardiac to test our hypothesis. Our investigation drew upon RNA-Seq gene expression data sourced from the BGEE database [41]. We assessed the degree of developmental co-expression among each mouse gene pair within the above-mentioned gene groups across five pivotal Theiler stages (TS17, TS19, TS21, TS23 and TS24) of mouse development, employing the Manhattan distance method (Fig 6). In this context, larger Manhattan distances indicate a greater divergence in expression between two genes.

**Fig 6:**
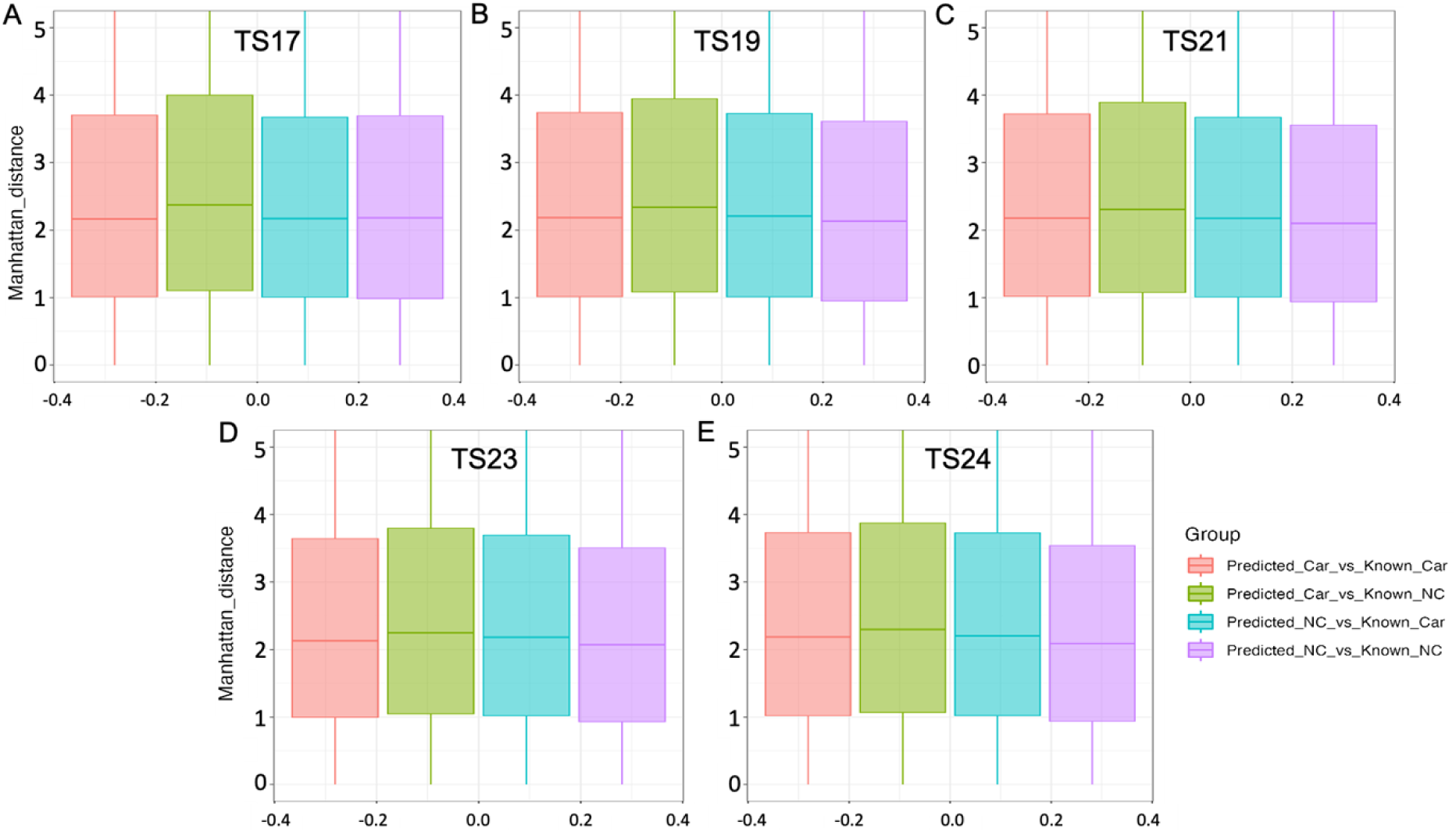
Measurements of co-expression. Differences in Manhattan across Theiler 17 (A), 19 (B), 21 (C), 23 (D) and 24 (E) stages in mice for four groups of gene pairs: (1) predicted cardiac vs known cardiac (Predicted_Car_vs_Known_Car), (2) predicted cardiac vs known non-cardiac (Predicted_Car_vs_Known_NC), (3) predicted non-cardiac vs known cardiac (Predicted_NC_vs_Known_Car), and (4) predicted non-cardiac vs known non-cardiac (Predicted_NC_vs_Known_NC). Here, ‘Manhattan_distance’ indicates the level of co-expression between gene pairs. Larger Manhattan distances signify a greater dissimilarity in expression between two genes.

Consistent with our expectations, we observed that predicted cardiac genes are more likely to exhibit similarities in developmental expression patterns with known cardiac genes in mouse (small distances and thus higher co–expression) throughout all Theiler stages when compared with both known non-cardiac genes and predicted non-cardiac genes, as confirmed by Kruskal-Wallis statistical tests (*P*-value < 0.05). Conversely, predicted non-cardiac genes demonstrated similar expression patterns with known non-cardiac genes, in comparison to known and predicted cardiac genes.

### Overlap with the ‘Unknome’

A recent study [42] has examined conserved genes of unknown function to determine if understudied genes have important biological roles. The authors have termed the collection of genes lacking functional annotations the ‘unknome’.

Experimental analysis of a subset of the unknome from *Drosophila* revealed most poorly annotated genes did indeed have attributable biological functions with clear phenotypes arising from mutations of these genes. We were therefore interested to determine if our predicted gene datasets contained genes within the unknome, and if our predictions might serve as a further way to assist with functional annotations of understudied genes. We found that 74% of the *Drosophila* unknome genes studied by Rocha et al., (2023) had orthologues in our predicted genes datasets (containing both cardiac and non-cardiac predictions). Given the yield of informative functional information arising from experimental study of the unknome in *Drosophila*, we propose that similar experimental investigation of our predicted gene datasets will identify novel biological associations. In particular, analysis of our predicted cardiac development gene dataset will be likely to identify pathways and molecules functioning in the process of heart development which are currently unknown.

### Comparison to New Cardiac Mouse Knockouts

As mentioned above, experimental analysis of our predictions is needed to confirm if our predicted cardiac gene dataset does indeed include genes newly found to play a role in cardiac development. To explore the experimental evidence, we conducted a search in the MGI database using the phenotype annotation ‘abnormal heart morphology’ to identify mouse knockouts with cardiac defects published after the generation of our classifier. Our aim was to obtain a set of genes that were not part of our training datasets, which now have experimentally determined cardiac development status. This search yielded a total of 68 genes (as of March 21, 2024). To assess the accuracy of our classifier, we compared these recently published cardiac genes with our gene predictions and observed a significant overlap with our predicted cardiac genes, as illustrated in Table S12. Notably, our classifier correctly predicted 78% (53 out of 68 genes) of these new cardiac knockouts to be associated with cardiac development, with false negatives tending to have lower confidence scores. The top 10 genes from this new dataset are listed in Table 7. This substantial alignment between our predictions and the new cardiac development genes underscores the efficacy of our classifier’s performance and validates the robustness of our approach in identifying genes involved in cardiac development.

**Table 7:** Top 10 recently published mouse cardiac genes and their prediction status by our Random Forest classifier.

| Gene Name | Encoded protein and function | Predicted Class | Confidence |
| --- | --- | --- | --- |
| Plvap | Plasmalemma vesicle-associated protein; involves in the formation of the diaphragms that bridge endothelial fenestrae | Cardiac | 0.97 |
| Smarca4 | Smarca4 protein; involves in transcriptional activation and repression | Cardiac | 0.945 |
| Pfkm | Phosphofructokinase, muscle; catalyses the phosphorylation of D-fructose 6-phosphate to fructose 1,6-bisphosphate by<br>ATP | Cardiac | 0.92 |
| Csf1 | Macrophage colony-stimulating factor 1; plays an essential role in the regulation of survival, proliferation and differentiation of hematopoietic precursor cells | Cardiac | 0.895 |
| Thbs1 | Thrombospondin-1; mediates cell-to-cell and cell-to-matrix interactions | Cardiac | 0.89 |
| Axl | AXL receptor tyrosine kinase; transduces signals from the extracellular matrix into the cytoplasm by binding growth factor GAS6 | Cardiac | 0.89 |
| Ager | Advanced glycosylation end product-specific receptor; senses endogenous stress signals with a broad ligand repertoire | Cardiac | 0.875 |
| Pnpla2 | Patatin-like phospholipase domain-containing protein; catalyses the initial step in triglyceride hydrolysis in adipocyte and non-adipocyte<br>lipid droplets | Cardiac | 0.86 |
| Mmp2 | 72 kDa type IV collagenase;<br>involves in angiogenesis, tissue<br>repair, tumor invasion,<br>inflammation | Cardiac | 0.86 |
| Thbs2 | Thrombospondin-2; mediates<br>cell-to-cell and cell-to-matrix<br>interactions | Cardiac | 0.85 |

### Zebrafish CRISPR knock down identifies a role for polr2h in heart development

Our predicted gene list included many *Polr* genes. *POLR1* genes have been implicated in various craniofacial disorders, and zebrafish *polr1a* and *polr1d* mutants exhibit severe craniofacial defects. Reports on zebrafish *polr2b* and *polr2d* have demonstrated roles in early zebrafish development, but roles in heart development have not been reported. Therefore, we wanted to test our prediction that *Polr2h* is needed during heart development by using CRISPR knock down in zebrafish embryos to evaluate organismal phenotype. The zebrafish genome has a single *POLR2H* ortholog, *polr2h*, but its role in zebrafish development has not yet been studied [43]. To test the function of zebrafish *polr2h*, we turned to an efficient CRISPR knock down approach, using a pool of four CRISPR guide RNAs [44].

Knock-down of polr2h caused embryos to have very characteristic small heads and heart that fail to loop properly at 48 hours post fertilization (hpf; Fig 7A-D).

**Figure 7.**
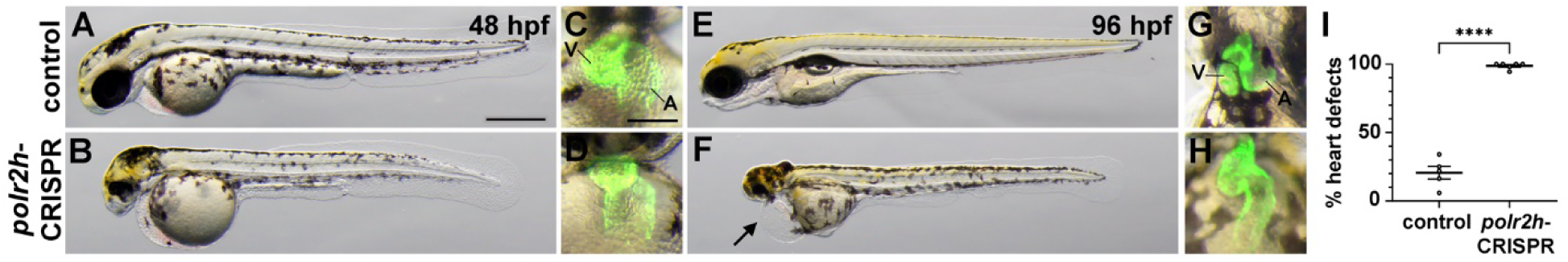
*polr2h* CRISPR knock down in zebrafish causes embryonic heart defects. A.-B. Lateral views of 48 hpf CRISPR control embryo (A) and *polr2h*-CRISPR embryo (B). C.-D. Ventral views of 48 hpf embryos shown in A. and B., respectively, showing *myl7-gfp*-expressing hearts in green. V, ventricle. A, atrium. E.-F. Lateral views of 96 hpf CRISPR control embryo (E) and *polr2h*-CRISPR embryo (F). Arrow in (F) points to cardiac oedema. G.-H. Ventral views of 96 hpf embryos shown in E. and F., respectively, showing *myl7-gfp*-expressing hearts in green. V, ventricle. A, atrium. Scale bars in A, C=500 µm. I. Graph showing frequency of heart defects observed in *polr2h*-CRISPR embryos at 96 hpf. Each dot represents an experimental replicate batch of embryos. N=5 replicates per condition. Each replicate batch consists of 17-54 embryos. **** *P*-value <0.0001, using 2-tailed t-test.

By 96 hpf, *polr2h-*CRISPR embryos continue to have very reduced heads, show oedema around the heart cavity, and have hearts in which the chambers are malformed and not properly looped (Fig7 E-H). Nearly 100% of *polr2h*-CRISPR embryos exhibit heart defects at 96 hpf (Fig7 I). The heart defects in these *polr2h* knock-down zebrafish embryos provide support for *POLR2H* as a candidate CHD gene in humans.

### Comparison to Human CHD Genes

We cross-referenced our predicted cardiac development gene dataset with multiple datasets of human CHD genes, aiming to identify which CHD genes are orthologues of our predicted cardiac genes. Several recent studies have highlighted genes associated with human CHDs [11, 13, 45–50]. The mouse orthologues of many of these CHD genes were not included in our machine learning training datasets. We observed a significant overlap between our cardiac development gene predictions and these newly identified CHD gene datasets (Table 8 and S13). Overall, 75% of the newly identified CHD genes are orthologues of mouse genes that we predicted to be associated with cardiac development, while the remaining genes are non-cardiac. The high percentage of overlap between our predictions and published CHD causative genes suggests that the cardiac development genes identified by our classifier are highly likely to be potential human CHD gene candidates.

**Table 8.**
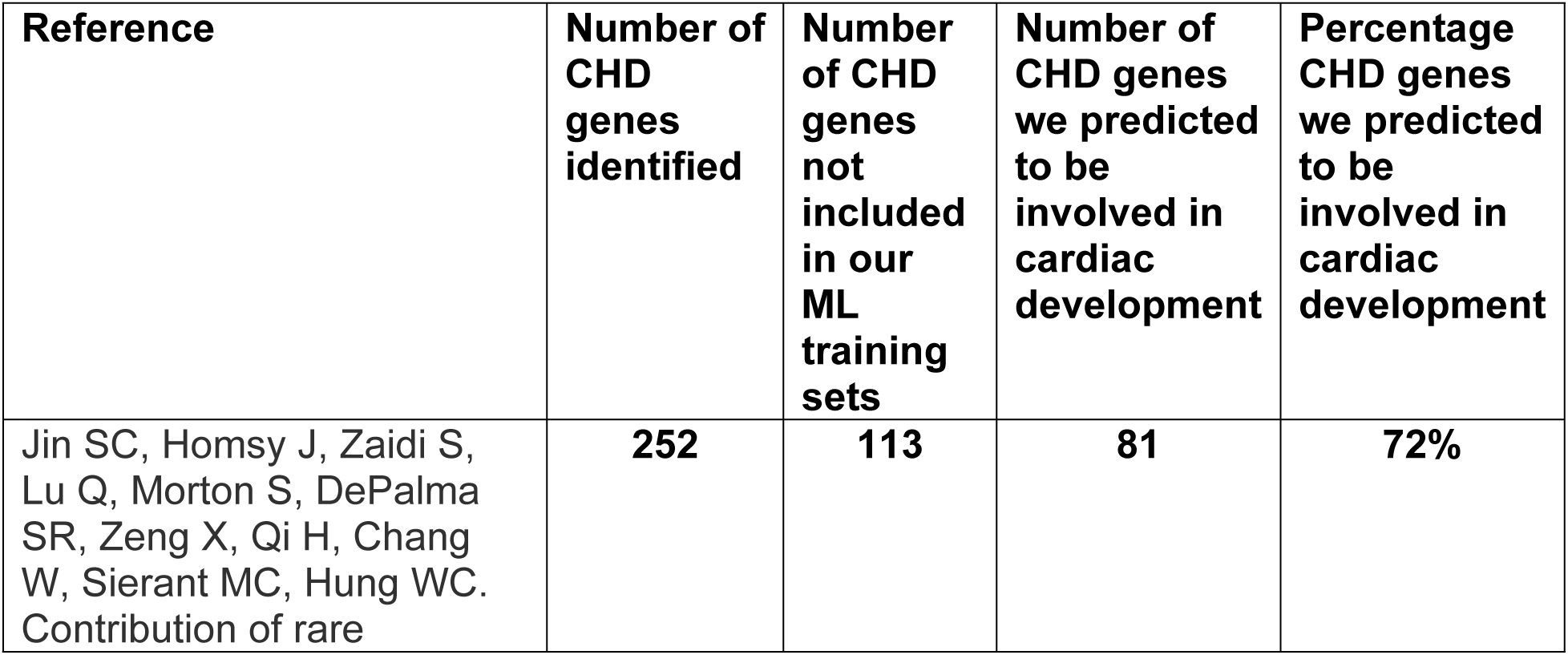
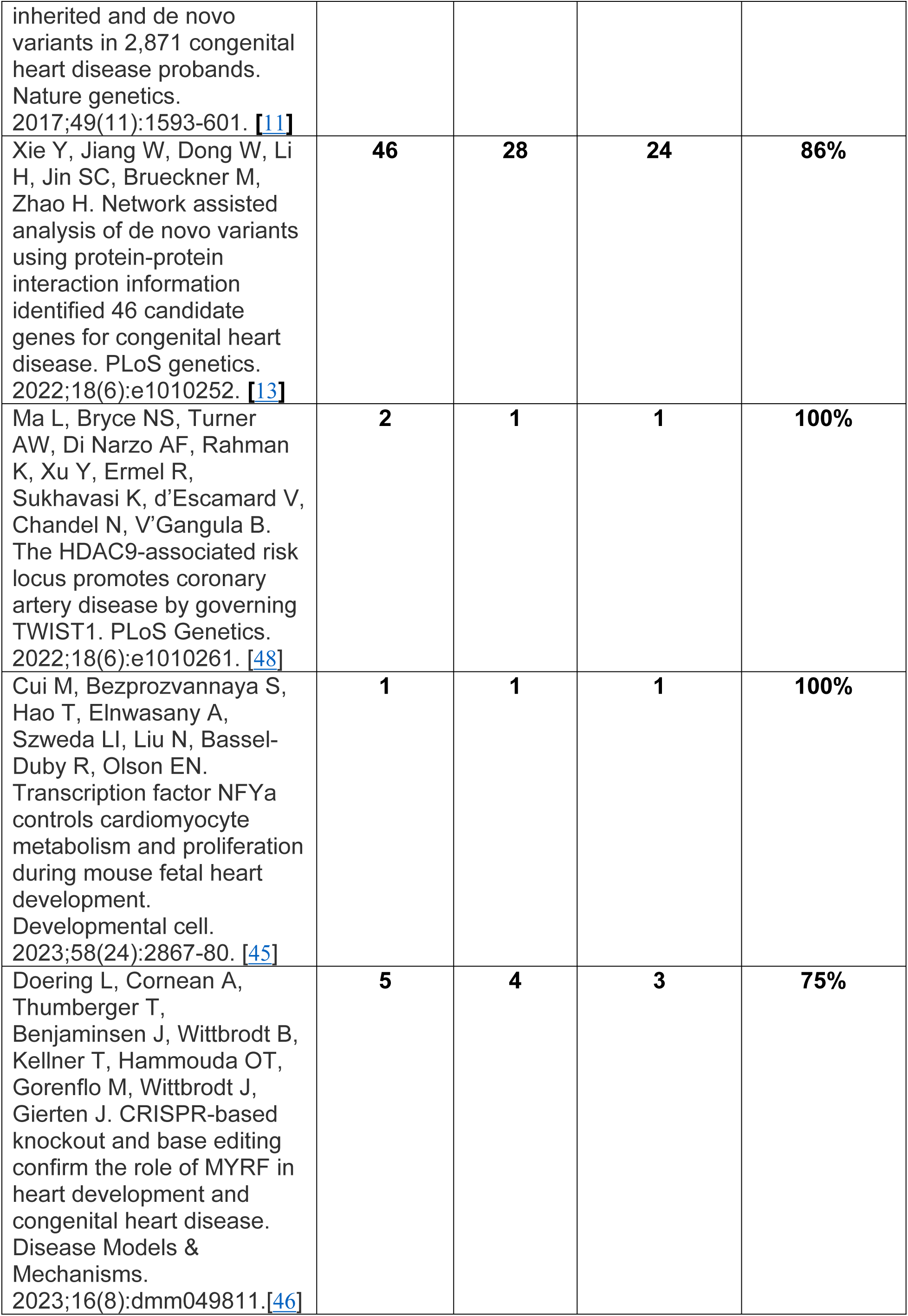
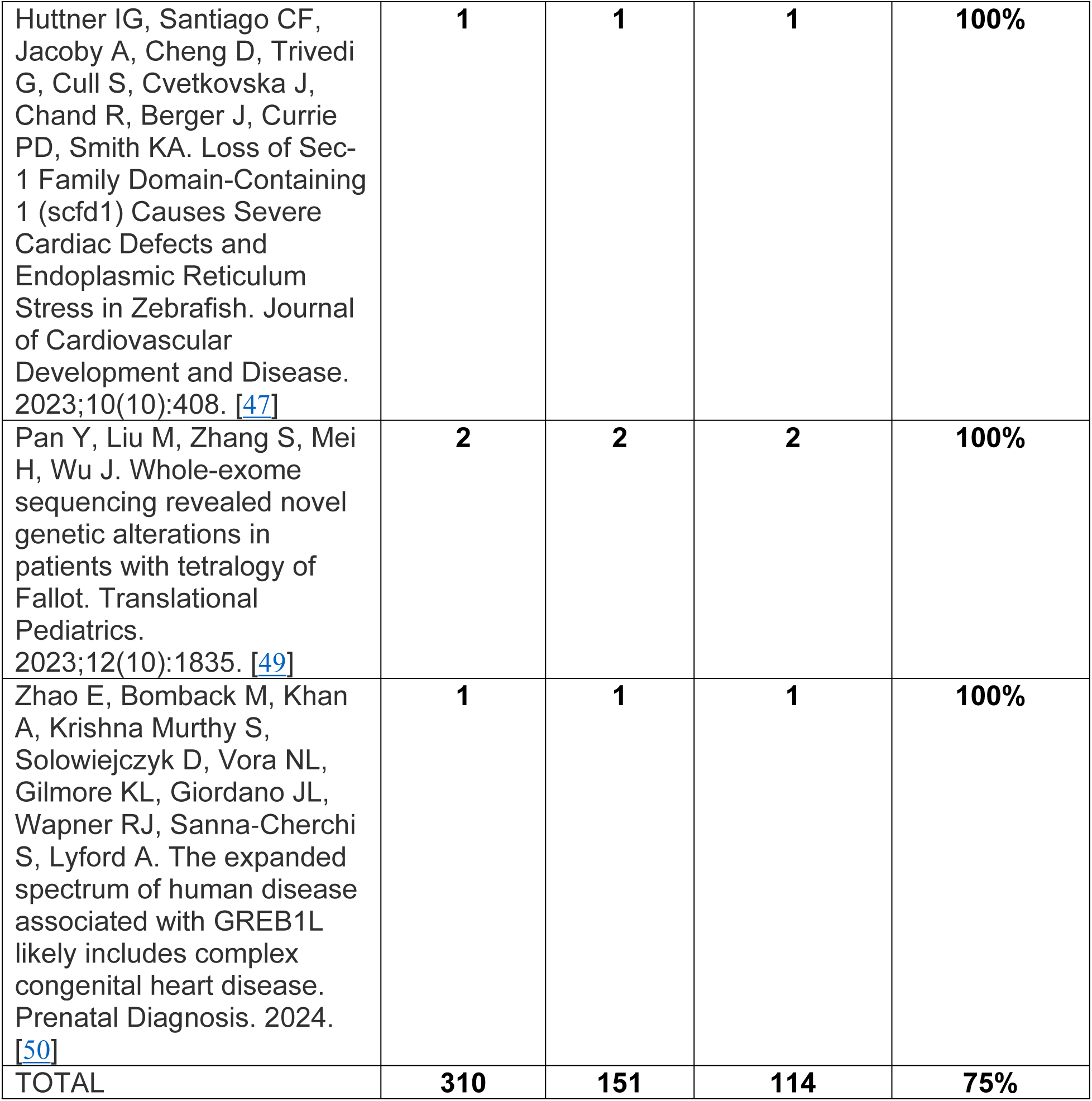
Cardiac development gene prediction matches to human CHDs. We compared the human orthologues of our mouse cardiac development gene predictions with human CHD genes identified in literature spanning from 2017 to 2024 to assess our classifier’s effectiveness in identifying CHD genes. If our classifier predicts a CHD gene as associated with cardiac development, we consider this prediction an accurate outcome.

We also assessed a total of 25 genes categorised as “Green” PanelApp [51] genes from the “Familial non-syndromic congenital heart disease” clinical testing panel (https://panelapp.genomicsengland.co.uk). These were genes included on the Genomics England panel used to assess participants with CHD recruited to the UK100KGP cohort and are therefore known human CHD disease genes. Our classifier correctly identified 88% (22 out of 25) of these genes as known or predicted cardiac genes, while the remaining 3 genes were classified as known or predicted non-cardiac genes.

We then evaluated a dataset containing 79 novel candidate genes found to contain high impact, rare, *de novo* variants identified in individual human CHD cases within the UK100KG cohort (Table S13). Of these, 47% were orthologues of predicted or known cardiac genes in our mouse datasets, while 20% were predicted non-cardiac genes. This demonstrates our classifier’s capability to recognise human CHD genes effectively. Table 9 displays the 10 highest-ranking candidate CHD genes, with the complete list available in Table S13.

**Table 9:** Top 10 classifier predictions for novel candidate CHD genes identified from the UK100KG cohort.

| Gene Name | Encoded Protein and Function | Predicted Class | Confidence |
| --- | --- | --- | --- |
| CSTF3 | Cleavage stimulation factor subunit 3; required for polyadenylation and 3'-end cleavage of mammalian pre-mRNAs | Cardiac | 0.955 |
| UBR2 | E3 ubiquitin-protein ligase UBR2; plays a critical role in chromatin inactivation | Cardiac | 0.955 |
| ERC1 | ELKS/Rab6-interacting/CAST family member 1; regulatory subunit of the IKK complex | Cardiac | 0.935 |
| CBLB | E3 ubiquitin-protein ligase<br>CBL-B; promotes ubiquitin degradation by the proteasome | Cardiac | 0.93 |
| PFKFB2 | 6-phosphofructo-2-kinase/fructose-2,6-bisphosphatase 3;<br>Catalyses both the synthesis and degradation of fructose 2,6-bisphosphate | Cardiac | 0.87 |
| NR4A1 | Nuclear receptor subfamily 4 group A member 1; orphan nuclear receptor | Cardiac | 0.865 |
| ABCF3 | ATP-binding cassette subfamily F member 3; displays an antiviral effect against flavivirus | Cardiac | 0.86 |
| AC008755.1 (ZC3H4) | Zinc finger CCCH domain-containing protein 4; RNA-binding protein that suppresses transcription of long non-coding RNAs | Cardiac | 0.85 |
| PYGB | Glycogen phosphorylase,<br>brain form; regulates<br>glycogen mobilization | Cardiac | 0.84 |
| CDC5L | Cell division cycle 5-like<br>protein; DNA-binding protein<br>involved in cell cycle control | Cardiac | 0.805 |

## Discussion

Advances in genomic research methodologies have expanded access to genome sequencing for CHD patients. However, a significant challenge arising from genomic analysis is linking variation within a specific gene to CHD causation in an individual patient. Machine learning, with its ability to uncover patterns within datasets, presents a promising method for identifying CHD-related genes. In this study, we developed a Random Forest (RF) classifier to identify genes likely required for cardiac development. These genes may serve as promising candidates for causing CHDs when mutated.

Our RF classifier achieved an 81% accuracy in a 10-fold cross-validation analysis following feature selection. The classifier demonstrated an 88% accuracy in predicting the cardiac development association status of known non-cardiac genes not utilised during the classifier’s training. Additionally, the classifier accurately predicted the cardiac development status for 88% of cardiac reporter line genes.

These results suggest that our classifier is not overfitted, as there is no drop in performance between training and testing. The high accuracy observed on these independent test datasets suggests that our classifier effectively discriminates between cardiac and non-cardiac genes. Furthermore, our predictions of the cardiac development status of all protein-coding genes in the mouse genome revealed that approximately 36% of genes may play a role in cardiac development, while 64% likely do not. In our training sets of genes with known cardiac development status, only 29% of the known phenotype genes have cardiac developmental defects, indicating that there are many cardiac development genes currently lacking experimental validation.

Subsequent examination of the protein network topology of the predicted cardiac and non-cardiac genes demonstrated that proteins within the predicted cardiac network exhibit a high degree of connectivity, aligning with our observation for the known cardiac genes. Additionally, the protein interaction networks for known cardiac genes and predicted cardiac genes have similar connectivity characteristics. Moreover, analysis of RNA-Seq gene expression data across development indicated that predicted cardiac genes have overlap with known cardiac genes in their developmental expression patterns. These findings support the hypothesis that the known and predicted cardiac genes jointly function in the process of cardiac development.

A large proportion of our predicted genes are orthologues of genes within the *Drosophila* unknome [42], which comprises genes without a current functional annotation. Our predictions can therefore begin to provide suggested functions for these genes, which may be borne out by further experimental analysis. Indeed, our predictions of genes associated with cardiac development do align with recently experimentally characterised mouse knockouts with abnormal cardiac development. Notably, our predictions matched 78% of the new cardiac knockouts, validating the robustness of our approach in identifying cardiac development genes. Additionally, we experimentally validated the prediction that *Polr2h* is needed for cardiac development through CRISPR knockdown in zebrafish. The phenotype seen in the *polr2h*-CRISPR embryos appears very similar to the published phenotypes for *polr2b* and *polr2d* [43, 52]. Maeta et al (2020) observed cardiac oedema in *polr2d* mutant embryos, although these previous studies did not directly examine heart development.

Comparison of the human orthologues of our predicted mouse cardiac genes with newly discovered human CHD genes revealed a 75% overlap, demonstrating the potential for our predictions to reveal candidate disease genes. One gene identified from newly reported mouse knockout experiments, *Smarca4* [11], has also recently been discovered to harbour pathological variants in patients with non-syndromic cleft lip and/or cleft palate and cardiac outflow tract defects [53]. An additional case report of a patient with Coffin-Siris syndrome with a novel variant in *SMARCA4* describes severe congenital heart disease as an aspect of the phenotype [54].

Our predicted cardiac genes may not fully overlap with CHD genes because our classifier was trained to identify genes involved in cardiac development, not on CHD gene features. Additionally, it is likely that some cardiac development genes will perform essential functions, and deleterious changes in those genes will cause embryonic lethality. Many of our known cardiac training set genes have embryonic lethal knockout phenotypes, confirming the essentiality of these genes.

Consequently, individuals harbouring deleterious variants in these genes may not survive development, and therefore these genes would not be identified from CHD patient cohorts. Nevertheless, our classifier identified large numbers of candidate genes that can be examined through future experimental analysis as potential causative genes for CHD.

Our ML approach offers a rapid, cost-effective, and complementary means of identifying cardiac developmental genes, serving as a valuable complement to experimental methods. Through a single analysis, our approach yielded cardiac predictions for the entire genome. The ranked list of genome wide cardiac developmental gene predictions, available in our publicly accessible database, will be of significant value to researchers and clinicians investigating genetic causes of CHDs, and could serve as a complement to the PanelApp traffic-light system [51]. By enabling prioritisation of genes during the evaluation of CHD patient sequence data, this resource has the potential to accelerate genetic diagnoses, thereby facilitating more informed treatment and disease management strategies for individuals affected by CHD. Further investigation of our predicted cardiac genes should elucidate their roles in cardiac development and reveal potential clinical associations with CHD.

## Materials and Methods

### Cardiac and Non-cardiac Mouse Gene Datasets

The MGI database was used to compile a dataset encompassing all mouse genes. To identify genes pertinent to cardiac development, we relied on information from the mouse knockout literature. This study included only null alleles of mouse genes, characterised by known phenotypes resulting from single gene knockout experiments (targeted deletions). Mouse genes were categorised as either cardiac or non-cardiac based on the mutant mouse phenotype data retrieved from the MGI database (as of June 17, 2021). We defined the phenotype of a knockout mouse as “cardiac” if the gene was known to be involved in cardiac development. These genes can potentially cause CHDs when mutated. Mouse knockouts with phenotypes known not to be associated with cardiac development were marked as “non-cardiac”. Some entries in these knockout datasets were ambiguous, annotated as both cardiac and non-cardiac in MGI. To resolve this ambiguity, we manually cross-referenced the phenotypes of these overlapping entries with published literature and included each gene in only a single category consistent with published phenotype descriptions. Mouse genes that were not labelled as either cardiac or non-cardiac were categorised as genes with “unknown” cardiac association status. Our datasets were limited to protein-coding genes only. We identified the encoded proteins for each mouse gene using the UniProt [55] (release 2021_3) database, selecting only the longest protein isoform for analysis per gene. Additionally, we obtained the Ensembl [56] gene identifier and UniGene [57, 58] expression cluster identifier mapping for each MGI gene symbol.

### Feature Collation

In machine learning, the individual pieces of information inputted into a classifier are termed ‘features’. These features are utilised by the classifier to recognise patterns that enable the prediction of whether a novel gene possesses similarities with genes within a specified training set. For our study, we assembled various features derived from gene and protein sequences, reflecting diverse aspects of mouse biology to distinguish between cardiac and non-cardiac phenotypes. These features encompass characteristics such as sequence properties, protein localisation and interaction details, developmental expression data, and gene ontology annotations (Table S14). Computation of features such as ‘gene length’, ‘GC content’, ‘transcript count’, ‘exon count’, ‘exon length’, and ‘intron length’ was performed using data extracted from the Ensembl database (release 103 of Mus musculus genes) through the Ensembl BioMart [59] data mining tool. Gene expression data, measured as transcripts per million (TPM), were sourced from the UniGene database for 13 embryonic developmental stages. Additionally, RNA-seq gene expression data spanning 8 tissue types (8 weeks heart, 8 weeks fibroblast, multi-cell lifecycle, zygote, Ths10 epiblast, 8 weeks thymus, 24 weeks adipose and stem cell) were obtained from the BGEE [41] v15.2 database. Protein characteristics including length, molecular weight, and amino acid composition were calculated using the Pepstats [60] program, with subcellular localisation features obtained from UniProt.

Evolutionary age, signal peptides, transmembrane domains, and subcellular locations were acquired from Ensembl, SignalP [61], and UniProt. Mouse protein-protein interaction (PPI) data were sourced from the I2D [24] v2.3 database, and various properties of PPI networks were computed using the ‘network analyzer’ plugin of Cytoscape [62] v3.9.1 and the Hubba web-based service. PPI data were also obtained from the BioGRID [38] v4.4 and STRING [39] v12.0 databases to conduct the PPI network analyses. Prior to network analyses, self-loops and duplicated edges were removed. Gene ontology (GO) terms were retrieved using the ‘Functional Annotation’ tool of the DAVID [27] v2021q4 web-based application.

Chromosome location data for all protein-coding mouse genes were obtained from Ensembl. Detailed explanations of these compiled features have already been provided in previous studies [32, 33, 63]. Furthermore, gene-level attributes, including gene constraint metrics such as probability of loss-of-function score (pLoF) and probability of being loss-of-function intolerant score (pLI), were obtained as features from the gnomAD v2.1.1 database [64].

### Machine Learning Classifiers

In this study, we developed a Random Forest (RF) classifier by including various features to discern genes highly likely to play a role in cardiac development, potentially serving as promising candidates for CHDs when mutated. We used WEKA v3.8.6, an openly accessible machine learning software, alongside R [65] for classifier implementation. To address the imbalanced classification issue, wherein a bias toward the larger gene group could occur, balanced training datasets were generated by employing random subsampling with no replacement [66] to ensure equal representation of cardiac and non-cardiac genes. Additionally, adjustments were made to refine features by replacing missing values with the respective feature mean values. Employing the 10-fold cross-validation technique on a dataset comprising both cardiac and non-cardiac mouse genes facilitated the prevention of overfitting; a classifier overfits when its prediction accuracy is high on the training dataset but poor on the test dataset. This approach entailed random partitioning of the training dataset into 10 equal subsets, with nine used for classifier training and one for testing. To classify a new gene, the features of the gene are evaluated using each of the decision trees within our Random Forest classifier. Each tree provides a class prediction or “vote,” and the class with the highest number of votes is chosen as the class to which the gene belongs. Our classifier produces a probability score that reflects the confidence level associated with prediction outcomes. This probability score is derived by averaging all predictions generated by decision trees within the Random Forest ensemble. Separate test datasets were generated from mouse genes not utilised in classifier training; evaluation of our classifier performance was assessed by estimating the proportion of accurately predicted genes within these test datasets.

### Feature Selection

The correctness of the classification depends significantly on the quality of the input features utilised in developing the classifier. Not all features within the training dataset are beneficial for this purpose. Irrelevant or redundant features can introduce noise into the classifier and lead to overfitting. Feature selection, therefore, is an imperative step for classification problems, especially when dealing with datasets containing numerous features. Its primary aim is to identify and select the most informative features relevant for classification, thus reducing overfitting and enhancing classification performance. Feature selection was performed employing the Information Gain filter method implemented in WEKA. This method estimates the significance of each feature by assessing its information gain with respect to the classification target [67]. Subsequently, only the most informative features were selected for classification, prioritised based on their significance. A higher information gain value indicates greater contribution of the feature in determining the classification target.

### Evaluation Metrices

The performance of our machine learning classifier was evaluated using various metrics, including recall, precision and accuracy (defined by equations (1) — (3)), where *TP*, *TN*, *FP*, and *FN* represent the number of true positives (i.e., correctly identified cardiac genes), true negatives (i.e., correctly identified non-cardiac genes), false positives (i.e., incorrectly identified non-cardiac genes), and false negatives (i.e., incorrectly identified of cardiac genes), respectively. Additionally, the area values of receiver operating characteristic curve (AUC) and precision-recall curve

(PRC) and confusion matrix were also considered for classifier evaluation. The ROC area provides a measure of overall classifier performance, while the PRC area evaluates the classifier’s performance in identifying samples from individual groups.

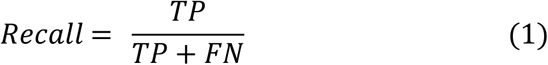

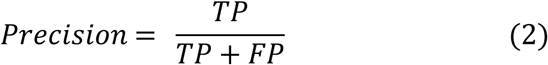

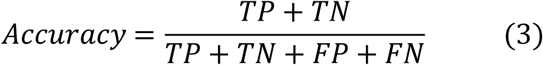

### Gene Co-expression Analysis

To examine if cardiac genes have similarities in developmental expression patterns, RNA-Seq gene expression data were obtained from the latest version of BGEE (release 15) which contains mouse expression data compiled from 566 libraries. We retrieved tissue-based RNA-Seq expression data represented as transcripts per million (TPM) for Theiler developmental stages 17, 19, 21, 23, and 24. These TPM values were transformed into corresponding log values using equation 4 to measure co-expression between every gene pair.

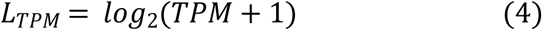

We categorised gene pairs into four groups: (1) predicted cardiac vs known cardiac, (2) predicted cardiac vs known non-cardiac, (3) predicted non-cardiac vs known cardiac, and (4) predicted non-cardiac vs known non-cardiac. We employed

Manhattan and Euclidean distance methods to compute numerical scores representing gene co-expression for each of these categories. These distance values were used to assess gene expression patterns between every gene pair across each Theiler stage. For two mouse genes a and b with expression values, the Manhattan distances between them were calculated using equation 5. Log TPM data (as per equation 4) were employed for the Manhattan distance calculation. Lower distances indicate higher co-expression between genes.

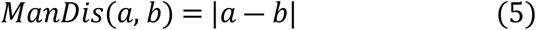

### Unknome Comparison

We retrieved mouse orthologues for a list of 358 *Drosophila* genes using the DRSC Integrative Ortholog Prediction Tool (DIOPT) [68]. The genes that lacked known functions formed the ‘unknome’ [42].

### RT-PCR verification of most confidently predicted cardiac genes

Embryonic and adult heart tissues were isolated from wild type 129S5/SvEvBrd mice and homogenised in Trizol. Total RNA was extracted, and complementary DNA (cDNA) was synthesised using High-Capacity cDNA Reverse Transcription Kit (Applied Biosystems). RT-PCR was conducted using PCR-Biosciences Ultra Mix Red polymerase with primers (Table S14) designed to amplify top predicted cardiac genes. PCR products were analysed by agarose gel electrophoresis and visualised under a UV transilluminator. 18S rRNA expression was used as a positive control for confirming the presence of cDNA template in all reactions. All animal experiments followed ethical guidelines at the Biological Service Facility, University of Manchester, UK (project licence number PP3720525) in compliance with Home Office regulations.

### Zebrafish husbandry and CRISPR

All experiments involving live zebrafish (*Danio rerio*) were carried out in compliance with Seattle Children’s Research Institute’s Institutional Animal Care and Use Committee guidelines. Zebrafish were raised and staged as previously described [69]. Time indicated as hpf refers to hours post-fertilization at 28.5°C.

The wild-type stock and genetic background used was AB. The *Tg(myl7:EGFP)^twu34^* line has been previously described [70]. For fish stock maintenance, eggs were collected from 20–30 min spawning periods and raised in Petri dishes in ICS water in a dark 28.5°C incubator, up to 5 dpf. After 5 dpf, the fish were maintained on a recirculating water system (Aquaneering) under a 14 h on, 10 h off light cycle. From 6–30 dpf, the fish were raised in 2.8 L tanks with a density of no more than 50 fish per tank and were fed a standard diet of paramecia (Carolina) one time per day and Zeigler AP100 dry larval diet two times per day. From 30 dpf onwards, the fish were raised in 6 L tanks with a density of no more than 50 fish per tank and were fed a standard diet of Artemia nauplii (Brine Shrimp Direct) and Zeigler adult zebrafish feed, each two times per day.

The sequences for the oligonucleotides (Table 10) used to synthesize the single-guide RNAs for *polr2h* CRISPR knockdown were taken from the genome-scale Lookup Table provided by [44]. For a negative control 4-guide set, we used the “Screen Scramble” guides of [44]. sgRNAs were synthesized from a pool of the four polr2h or control oligo as in [44]. Each oligo incorporates a T7 RNA polymerase site, annealed to a common scaffold oligo, as described in Wu et al. For phenotype analysis, 2 µL of a 4-guide cocktail of sgRNA at 2 µg/µL was combined with 2 µL of Cas9 protein (IDT Alt-R S.p. Cas9 Nuclease V3) at 10 µM (diluted as in [44]) and incubated at 37°C for 5 minutes. One microliter of phenol red injection solution (0.1% phenol red and 0.2M KCl in water) was added to generate the working solution for embryo injections. One-cell stage embryos, collected from *Tg(myl7:EGFP)^twu34^* fish, were injected in the yolk with one nanolitre of the RNP working solution. Following 1-cell stage injections, embryos were allowed to develop further and GFP expression was assessed in live embryos. For imaging, representative live embryos were anaesthetized in tricaine (Sigma) and then transferred to 2.5% methyl cellulose (Sigma) in ICS water. Images were captured with an Olympus DP74 camera mounted on an Olympus SZX16 stereomicroscope using cellSens Dimension imaging software. Merged brightfield and GFP fluorescence images were made in Adobe Photoshop.

**Table 10.**
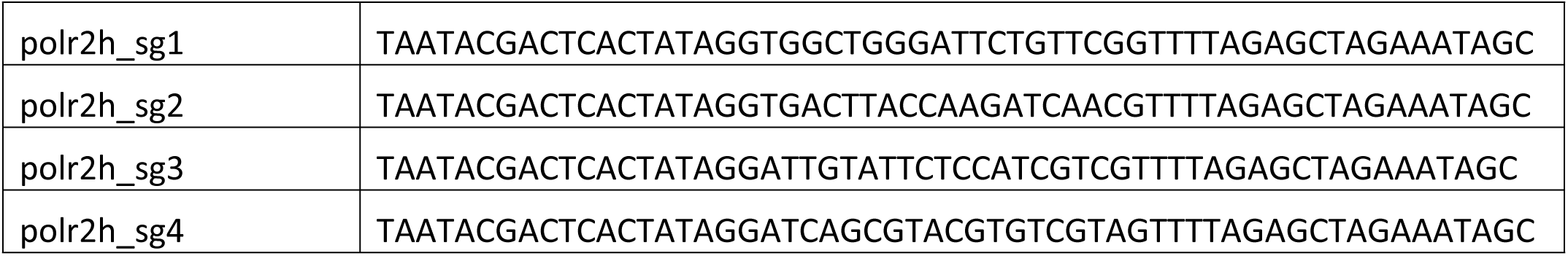
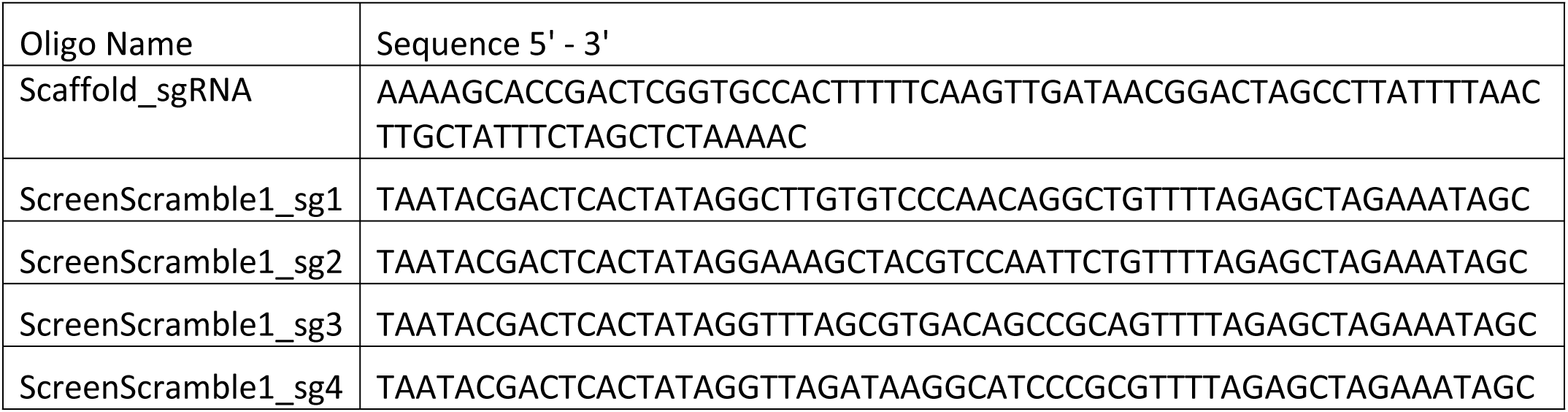
Oligonucleotides for sgRNA synthesis for G0 CRISPANT analysis.

### Statistics

Statistical analyses were conducted using SPSS v27 and R software packages. The significance of each feature was assessed using the non-parametric Mann–Whitney *U* test. Additionally, we employed the Chi-squared (χ²) test to examine whether the observed frequencies of a feature in the cardiac and non-cardiac datasets differed from expected values. The Bonferroni correction [71] was applied to correct the p–values for multiple testing.

## Acknowledgments

This research was made possible through access to data in the National Genomic Research Library, which is managed by Genomics England Limited (a wholly owned company of the Department of Health and Social Care). The National Genomic Research Library holds data provided by patients and collected by the NHS as part of their care and data collected as part of their participation in research. The National Genomic Research Library is funded by the National Institute for Health Research and NHS England. The Wellcome Trust, Cancer Research UK and the Medical Research Council have also funded research infrastructure.

## Funding Statement

This research was supported by British Heart Foundation grant PG/20/14/35030 to AJD and KEH and PG/22/11127 to KEH and DT, BHF project grant PG/18/28/33632 to CAJ and KH, MRC Clinical-Academic Research Partnership (CARP) fellowship MR/V037617/1 to VH, and Additional Ventures Foundation Single Ventricle Research Fund (SVRF) award and Department of Defense/USAMRAA grant #W81XWH2010433 to LM. BK is supported by a BHF personal chair.

## Notes

### Competing Interest Statement

The authors have declared no competing interest.

## References

1. Fahed AC, Gelb BD, Seidman J, Seidman CE. Genetics of congenital heart disease: the glass half empty. Circulation research. 2013;112(4):707–20.

2. Boneva RS, Botto LD, Moore CA, Yang Q, Correa A, Erickson JD. Mortality associated with congenital heart defects in the United States: trends and racial disparities, 1979–1997. Circulation. 2001;103(19):2376–81.

3. Zaidi S, Brueckner M. Genetics and genomics of congenital heart disease. Circulation research. 2017;120(6):923–40.

4. Khanna AD, Duca LM, Kay JD, Shore J, Kelly SL, Crume T. Prevalence of mental illness in adolescents and adults with congenital heart disease from the Colorado Congenital Heart Defect Surveillance System. The American journal of cardiology. 2019;124(4):618–26.

5. Bonthrone AF, Chew A, Kelly CJ, Almedom L, Simpson J, Victor S, et al. Cognitive function in toddlers with congenital heart disease: The impact of a stimulating home environment. Infancy. 2021;26(1):184–99.

6. Mandalenakis Z, Karazisi C, Skoglund K, Rosengren A, Lappas G, Eriksson P, et al. Risk of cancer among children and young adults with congenital heart disease compared with healthy controls. JAMA network open. 2019;2(7):e196762-e.

7. Gelb BD, Chung WK. Complex genetics and the etiology of human congenital heart disease. Cold Spring Harbor perspectives in medicine. 2014;4(7):a013953.

8. Blue GM, Kirk EP, Giannoulatou E, Dunwoodie SL, Ho JW, Hilton DC, et al. Targeted next-generation sequencing identifies pathogenic variants in familial congenital heart disease. Journal of the American College of Cardiology. 2014;64(23):2498–506.

9. Akhirome E, Walton NA, Nogee JM, Jay PY. The complex genetic basis of congenital heart defects. Circulation Journal. 2017;81(5):629–34.

10. Priest JR, Osoegawa K, Mohammed N, Nanda V, Kundu R, Schultz K, et al. De novo and rare variants at multiple loci support the oligogenic origins of atrioventricular septal heart defects. PLoS genetics. 2016;12(4):e1005963.

11. Jin SC, Homsy J, Zaidi S, Lu Q, Morton S, DePalma SR, et al. Contribution of rare inherited and de novo variants in 2,871 congenital heart disease probands. Nature genetics. 2017;49(11):1593–601.

12. Gordon DM, Cunningham D, Zender G, Lawrence PJ, Penaloza JS, Lin H, et al. Exome sequencing in multiplex families with left-sided cardiac defects has high yield for disease gene discovery. PLoS Genetics. 2022;18(6):e1010236.

13. Xie Y, Jiang W, Dong W, Li H, Jin SC, Brueckner M, et al. Network assisted analysis of de novo variants using protein-protein interaction information identified 46 candidate genes for congenital heart disease. PLoS genetics. 2022;18(6):e1010252.

14. Meehan TF, Conte N, West DB, Jacobsen JO, Mason J, Warren J, et al. Disease model discovery from 3,328 gene knockouts by The International Mouse Phenotyping Consortium. Nature genetics. 2017;49(8):1231–8.

15. Radivojac P, Clark WT, Oron TR, Schnoes AM, Wittkop T, Sokolov A, et al. A large-scale evaluation of computational protein function prediction. Nature methods. 2013;10(3):221–7.

16. Jiang Y, Oron TR, Clark WT, Bankapur AR, D’Andrea D, Lepore R, et al. An expanded evaluation of protein function prediction methods shows an improvement in accuracy. Genome biology. 2016;17:1–19.

17. Bult CJ, Eppig JT, Kadin JA, Richardson JE, Blake JA, Group MGD. The Mouse Genome Database (MGD): mouse biology and model systems. Nucleic acids research. 2008;36(suppl_1):D724–D8.

18. Aibar S, Fontanillo C, Droste C, De Las Rivas J. Functional Gene Networks: R/Bioc package to generate and analyse gene networks derived from functional enrichment and clustering. Bioinformatics. 2015;31(10):1686–8.

19. Yepes S, Tucker MA, Koka H, Xiao Y, Jones K, Vogt A, et al. Using whole-exome sequencing and protein interaction networks to prioritize candidate genes for germline cutaneous melanoma susceptibility. Scientific Reports. 2020;10(1):17198.

20. Siitonen A, Kytövuori L, Nalls MA, Gibbs R, Hernandez DG, Ylikotila P, et al. Finnish Parkinson’s disease study integrating protein-protein interaction network data with exome sequencing analysis. Scientific Reports. 2019;9(1):18865.

21. Sun PG, Gao L, Han S. Prediction of human disease-related gene clusters by clustering analysis. International journal of biological sciences. 2011;7(1):61.

22. Smedley D, Köhler S, Czeschik JC, Amberger J, Bocchini C, Hamosh A, et al. Walking the interactome for candidate prioritization in exome sequencing studies of Mendelian diseases. Bioinformatics. 2014;30(22):3215–22.

23. Ghiassian SD, Menche J, Barabási A-L. A DIseAse MOdule Detection (DIAMOnD) algorithm derived from a systematic analysis of connectivity patterns of disease proteins in the human interactome. PLoS computational biology. 2015;11(4):e1004120.

24. Brown KR, Jurisica I. Unequal evolutionary conservation of human protein interactions in interologous networks. Genome biology. 2007;8:1–11.

25. Lin C-Y, Chin C-H, Wu H-H, Chen S-H, Ho C-W, Ko M-T. Hubba: hub objects analyzer—a framework of interactome hubs identification for network biology. Nucleic acids research. 2008;36(suppl_2):W438–W43.

26. Ashburner M, Ball CA, Blake JA, Botstein D, Butler H, Cherry JM, et al. Gene ontology: tool for the unification of biology. Nature genetics. 2000;25(1):25–9.

27. Huang DW, Sherman BT, Tan Q, Kir J, Liu D, Bryant D, et al. DAVID Bioinformatics Resources: expanded annotation database and novel algorithms to better extract biology from large gene lists. Nucleic acids research. 2007;35(suppl_2):W169–W75.

28. He H, Garcia EA. Learning from imbalanced data. IEEE Transactions on knowledge and data engineering. 2009;21(9):1263–84.

29. Visa S, Ralescu A, editors. Issues in mining imbalanced data sets-a review paper. Proceedings of the sixteen midwest artificial intelligence and cognitive science conference; 2005: sn.

30. Breiman L. Random forests. Machine learning. 2001;45:5–32.

31. Hall M, Frank E, Holmes G, Pfahringer B, Reutemann P, Witten IH. The WEKA data mining software: an update. ACM SIGKDD explorations newsletter. 2009;11(1):10–8.

32. Kabir M, Stuart HM, Lopes FM, Fotiou E, Keavney B, Doig AJ, et al. Predicting congenital renal tract malformation genes using machine learning. Scientific Reports. 2023;13(1):13204.

33. Tian D, Wenlock S, Kabir M, Tzotzos G, Doig AJ, Hentges KE. Identifying mouse developmental essential genes using machine learning. Disease Models & Mechanisms. 2018;11(12):dmm034546.

34. Acencio ML, Lemke N. Towards the prediction of essential genes by integration of network topology, cellular localization and biological process information. BMC bioinformatics. 2009;10:1–18.

35. Bureau A, Dupuis J, Falls K, Lunetta KL, Hayward B, Keith TP, et al. Identifying SNPs predictive of phenotype using random forests. Genetic Epidemiology: The Official Publication of the International Genetic Epidemiology Society. 2005;28(2):171–82.

36. Yuan Y, Xu Y, Xu J, Ball RL, Liang H. Predicting the lethal phenotype of the knockout mouse by integrating comprehensive genomic data. Bioinformatics. 2012;28(9):1246–52.

37. Motenko H, Neuhauser SB, O’keefe M, Richardson JE. MouseMine: a new data warehouse for MGI. Mammalian Genome. 2015;26:325–30.

38. Stark C, Breitkreutz B-J, Reguly T, Boucher L, Breitkreutz A, Tyers M. BioGRID: a general repository for interaction datasets. Nucleic acids research. 2006;34(suppl_1):D535–D9.

39. Mering Cv, Huynen M, Jaeggi D, Schmidt S, Bork P, Snel B. STRING: a database of predicted functional associations between proteins. Nucleic acids research. 2003;31(1):258–61.

40. Tian W, Zhang LV, Taşan M, Gibbons FD, King OD, Park J, et al. Combining guilt-by-association and guilt-by-profiling to predict Saccharomyces cerevisiae gene function. Genome biology. 2008;9:1–21.

41. Bastian F, Parmentier G, Roux J, Moretti S, Laudet V, Robinson-Rechavi M, editors. Bgee: integrating and comparing heterogeneous transcriptome data among species. Data Integration in the Life Sciences: 5th International Workshop, DILS 2008, Evry, France, June 25-27, 2008 Proceedings 5; 2008: Springer.

42. Rocha JJ, Jayaram SA, Stevens TJ, Muschalik N, Shah RD, Emran S, et al. Functional unknomics: Systematic screening of conserved genes of unknown function. PLoS biology. 2023;21(8):e3002222.

43. Bradford YM, Van Slyke CE, Ruzicka L, Singer A, Eagle A, Fashena D, et al. Zebrafish information network, the knowledgebase for Danio rerio research. Genetics. 2022;220(4):iyac016.

44. Wu RS, Lam II, Clay H, Duong DN, Deo RC, Coughlin SR. A rapid method for directed gene knockout for screening in G0 zebrafish. Developmental cell. 2018;46(1):112–25. e4.

45. Cui M, Bezprozvannaya S, Hao T, Elnwasany A, Szweda LI, Liu N, et al. Transcription factor NFYa controls cardiomyocyte metabolism and proliferation during mouse fetal heart development. Developmental cell. 2023;58(24):2867–80. e7.

46. Doering L, Cornean A, Thumberger T, Benjaminsen J, Wittbrodt B, Kellner T, et al. CRISPR-based knockout and base editing confirm the role of MYRF in heart development and congenital heart disease. Disease Models & Mechanisms. 2023;16(8):dmm049811.

47. Huttner IG, Santiago CF, Jacoby A, Cheng D, Trivedi G, Cull S, et al. Loss of Sec-1 Family Domain-Containing 1 (scfd1) Causes Severe Cardiac Defects and Endoplasmic Reticulum Stress in Zebrafish. Journal of Cardiovascular Development and Disease. 2023;10(10):408.

48. Ma L, Bryce NS, Turner AW, Di Narzo AF, Rahman K, Xu Y, et al. The HDAC9-associated risk locus promotes coronary artery disease by governing TWIST1. PLoS Genetics. 2022;18(6):e1010261.

49. Pan Y, Liu M, Zhang S, Mei H, Wu J. Whole-exome sequencing revealed novel genetic alterations in patients with tetralogy of Fallot. Translational Pediatrics. 2023;12(10):1835.

50. Zhao E, Bomback M, Khan A, Krishna Murthy S, Solowiejczyk D, Vora NL, et al. The expanded spectrum of human disease associated with GREB1L likely includes complex congenital heart disease. Prenatal Diagnosis. 2024.

51. Martin AR, Williams E, Foulger RE, Leigh S, Daugherty LC, Niblock O, et al. PanelApp crowdsources expert knowledge to establish consensus diagnostic gene panels. Nature genetics. 2019;51(11):1560–5.

52. Maeta M, Kataoka M, Nishiya Y, Ogino K, Kashima M, Hirata H. RNA polymerase II subunit D is essential for zebrafish development. Scientific Reports. 2020;10(1):13213.

53. Munabi NC, Mikhail S, Toubat O, Webb M, Auslander A, Sanchez-Lara PA, et al. High prevalence of deleterious mutations in concomitant nonsyndromic cleft and outflow tract heart defects. American Journal of Medical Genetics Part A. 2022;188(7):2082–95.

54. Dsouza NR, Zimmermann MT, Geddes GC. A case of Coffin–Siris syndrome with severe congenital heart disease and a novel SMARCA4 variant. Molecular Case Studies. 2019;5(3):a003962.

55. Consortium U. UniProt: a hub for protein information. Nucleic acids research. 2015;43(D1):D204–D12.

56. Hubbard T, Barker D, Birney E, Cameron G, Chen Y, Clark L, et al. The Ensembl genome database project. Nucleic acids research. 2002;30(1):38–41.

57. Pontius JU, Wagner L, Schuler GD. 21. UniGene: A unified view of the transcriptome. The NCBI Handbook Bethesda, MD: National Library of Medicine (US), NCBI. 2003.

58. Stanton J-AL, Macgregor AB, Green DP. Identifying tissue-enriched gene expression in mouse tissues using the NIH UniGene database. Applied bioinformatics. 2003;2:S65–S74.

59. Smedley D, Haider S, Ballester B, Holland R, London D, Thorisson G, et al. BioMart–biological queries made easy. BMC genomics. 2009;10(1):1–12.

60. Rice P, Longden I, Bleasby A. EMBOSS: the European molecular biology open software suite. Trends in genetics. 2000;16(6):276–7.

61. Petersen TN, Brunak S, Von Heijne G, Nielsen H. SignalP 4.0: discriminating signal peptides from transmembrane regions. Nature methods. 2011;8(10):785–6.

62. Shannon P, Markiel A, Ozier O, Baliga NS, Wang JT, Ramage D, et al. Cytoscape: a software environment for integrated models of biomolecular interaction networks. Genome research. 2003;13(11):2498–504.

63. Kabir M, Barradas A, Tzotzos GT, Hentges KE, Doig AJ. Properties of genes essential for mouse development. PLoS One. 2017;12(5):e0178273.

64. Karczewski KJ, Francioli LC, Tiao G, Cummings BB, Alföldi J, Wang Q, et al. The mutational constraint spectrum quantified from variation in 141,456 humans. Nature. 2020;581(7809):434-43.

65. R Core Team R. R: A language and environment for statistical computing. 2013.

66. Vitter JS. Random sampling with a reservoir. ACM Transactions on Mathematical Software (TOMS). 1985;11(1):37–57.

67. Han J, Pei J, Tong H. Data mining: concepts and techniques: Morgan kaufmann; 2022.

68. Hu Y, Flockhart I, Vinayagam A, Bergwitz C, Berger B, Perrimon N, et al. An integrative approach to ortholog prediction for disease-focused and other functional studies. BMC bioinformatics. 2011;12:1–16.

69. Westerfield M. The Zebrafish Book; A guide for the laboratory use of zebrafish (Danio rerio). (No Title). 2007.

70. Huang CJ, Tu CT, Hsiao CD, Hsieh FJ, Tsai HJ. Germ-line transmission of a myocardium-specific GFP transgene reveals critical regulatory elements in the cardiac myosin light chain 2 promoter of zebrafish. Developmental dynamics: an official publication of the American Association of Anatomists. 2003;228(1):30–40.

71. Sedgwick P. Multiple significance tests: the Bonferroni correction. Bmj. 2012;344.

